# Multilayered Defense Responses in Sugarcane Against *Pratylenchus zeae* Revealed by Comparative Transcriptomics

**DOI:** 10.1101/2025.08.25.672178

**Authors:** PMS Confort, TG Souto, J Ferreira, S Creste, CB Monteiro-Vitorello

**Affiliations:** Departamento de Genética, Universidade de São Paulo, Escola Superior de Agricultura Luiz de Queiroz, Esalq, Piracicaba, SP, Brasil; Programa de Pós-Graduação em Fitopatologia, Departamento de Fitopatologia e Nematologia, Universidade de São Paulo, Escola Superior de Agricultura Luiz de Queiroz, Esalq, Piracicaba, SP, Brasil; Centro de Cana, Instituto Agronômico de Campinas, Ribeirão Preto, SP, Brasil

**Keywords:** Sugarcane, *Pratylenchus zeae*, Comparative transcriptomics, Plant–nematode interaction

## Abstract

The root-lesion nematode *Pratylenchus zeae* ranks among the most pervasive soilborne threats to global sugarcane (Saccharum spp.) production, however, the molecular basis of host resistance to this pathogen remains largely unexplored. Using comparative transcriptomics 15 days after inoculation, we profiled root response patterns of a resistant cultivar (RB966982) and a susceptible cultivar (CTC9001). Differential gene expression analysis identified 3,385 DEGs (1,426 up, 1,959 down) in the susceptible CTC9001 and 8,689 DEGs (5,334 up, 3,355 down) in the resistant RB966982 when comparing control and inoculated plants, revealing distinct genotype-specific defense strategies. The resistant genotype mounted a coordinated, multilayered defense marked by dramatic transcriptional reprogramming, including intensified glycolysis and fatty-acid biosynthesis, elevated oxidoreductase/ROS activity, and concurrent activation of jasmonic and salicylic acid associated pathways, with strong induction of multiple PR1 homologs. RB966982 also showed pre-primed and inducible resistance gene analogs and targeted cell-wall reinforcement through xyloglucan fucosylation. By contrast, CTC9001 displayed a delayed or attenuated response characterized by cell-proliferation signatures and broad activation of callose-related defenses (1,3-β-D-glucan synthesis) that appear insufficient to limit nematode progression. GO enrichment analysis revealed that the resistant response was dominated by biological processes linked to carbohydrate and lipid metabolism, oxidoreductase activity, hormone signaling, and cell-wall organization, while the susceptible response was enriched in stress-related pathways lacking coordinated metabolic and structural reinforcement. Collectively, these findings indicate that durable resistance to *P. zeae* is unlikely to arise from a single mechanism, effective protection will require stacking complementary defense layers, including early metabolic reprogramming, robust hormone-mediated signaling, and reinforced cell-wall barriers. The candidate RGAs, PR1 homologs, and cell-wall remodeling enzymes identified here, together with the GO-enriched pathways, provide concrete targets for marker-assisted breeding and gene editing strategies aimed at developing sugarcane cultivars with durable nematode resistance.

## 2. INTRODUCTION

Over 80% of global sugar production relies on the cultivation of sugarcane, a Poaceae species belonging to the *Saccharum* genus (Lakshmanan et al., 2005). In addition to sugar production, sugarcane serves as a key raw material for bioethanol, a renewable energy source of global relevance as the world races for sustainable fuels and toward reducing the ecological impacts caused by human activities (Goldemberg et al., 2008). Amid these efforts, Brazil has emerged as a global leader over the past half-decade, driven by substantial investments in scientific research focused on sugarcane and its agroindustrial complex (Lopes et al., 2016; Matsuoka et al., 2009).

The 2024/2025 Brazilian sugarcane harvest is expected to reach a volume of 678.67 million tons. This represents a reduction of 8.8% in productivity per hectare compared to the 2023/2024 harvest, despite a cultivation area 4.7% higher than the previous year. This decrease in productivity is related to lower rainfall and elevated temperatures in key production zones (CONAB, Companhia Nacional de Abastecimento 2024). In addition to these abiotic stresses, emerging diseases such as sugarcane wilting syndrome, caused by a complex of species including *Colletotrichum falcatum* and *Fusarium* spp., have been reported as one of the main driving forces behind this lower productivity in the current year (CONAB; IAC - Instituto Agronômico de Campinas, 2024). The average production for the current harvest in Brazil is 78 tons per hectare.

Globally, it is estimated that 10–20% of sugarcane production is lost due to the action of harmful organisms. As a high-yielding crop, even fractional percentage reductions in productivity can lead to immense losses (Huang et al., 2018). Among the pathogens capable of parasitizing sugarcane, soilborne pathogens pose a unique challenge for crop growers and researchers alike, given the perennial nature of this grass species. Sugarcane crop cycles starting with a planted cane can be followed by 4–6 ratoon crops, providing a stable substrate for root pathogens for several years (Mehnaz, 2013; Ren et al., 2024).

Among these soilborne pathogens, a total of 317 species of nematodes are known to parasitize sugarcane, accounting for 48 genera with both ecto-and endoparasitic behaviors. Among them, the *Pratylenchus* and *Meloidogyne* genera are two of the most commonly associated with sugarcane yield losses, with *Pratylenchus zeae* being the most prevalent species worldwide (Ramouthar & Bhuiyan, 2018).

*P. zeae* is a root-lesion nematode. Species of this genus are endoparasites that inhabit the root cortex, moving through and between parenchyma cells. This migration leads to the formation of necrotic regions, which appear as small lesions on washed roots. During migration, the nematode combines mechanical force from continuous stylet thrusting with the secretion of cell wall-degrading enzymes. Once the cell wall is breached, another set of enzymes is secreted into the cytoplasm before the cellular contents are ingested by the nematode (Castillo & Vovlas, 2010; Duncan & Moens, 2006). On sugarcane, *P. zeae* symptoms include red, reddish-purple, or brown lesions on roots. Depending on the timing of infection, lesions tend to become necrotic, causing the entire root system to darken. Above ground, reduced shoot number, leaf yellowing, and stalk shortening can usually be observed in patches of stunted plants (Ramouthar & Bhuiyan, 2018). Additionally, *P. zeae* presence is associated with a severe reduction in fine root hairs on sugarcane (Blair, 2005). These root structures play a key role in nutrient acquisition, water uptake, microbial interactions, and even root– soil cohesion, decreasing potential erosion (De Baets et al., 2020; Matsuoka & Garcia, 2011; Novero et al., 2009).

High-throughput sequencing technologies have reshaped research in plant and microbial biology, enabling a deeper understanding of genetic diversity within agricultural ecosystems (Adams et al., 2018; Nilsson et al., 2019). This is particularly true for plant pathogen genomes, which, once sequenced, serve as blueprints for uncovering potential virulence and avirulence factors, facilitating the prediction of genes associated with diverse pathogenic strategies (Aylward et al., 2017). Genome assemblies for nematodes had an early start, with *Caenorhabditis elegans* being the first multicellular eukaryote to have its genome sequenced 25 years ago, becoming a model organism for animal studies since then (Wilson, 1999). Despite this head start, genomic studies of phytonematodes lag behind those of other plant pathogens, especially in the genus *Pratylenchus*, which as of 2024 had only two sequenced genomes: *Pratylenchus scribneri* and *Pratylenchus coffeae* (Arora et al., 2023; Burke et al., 2015). The lack of genomic data for this genus presents an even greater challenge for studies in Brazil, where the most common species are *P. brachyurus* and *P. zeae* (Martinha et al., 2022; Santana-Gomes et al., 2019). For *Pratylenchus* spp., transcriptomic studies are just as scarce as genome assemblies, with most current research focused on the *Meloidogyne* and *Heterodera* genera. This can be partly attributed to the high economic impact of these two genera on crop yields (Jones et al., 2013; Opperman et al., 2009). However, economic relevance is not the only factor: assessing host resistance to root-lesion nematodes can be complicated, as visual evaluation of root and aboveground symptoms does not always reflect the pathogen’s actual reproductive capabilities, especially in C4 plants with vigorous root systems (Dababat & Fourie, 2018; Waele & Elsen, 2002).

Phenotyping is an essential step in evaluating host resistance to pathogens. Among the main methodologies used in the study of plant-parasitic nematodes, the reproduction factor (RF = PF/PI) is widely applied (Oostenbrink, 1966). This parameter measures the ratio between final and initial nematode populations and is considered one of the most reliable indicators for assessing cultivar resistance. Genotypes with RF values less than 1 are considered resistant, while those with RF values equal to or greater than 1 are classified as susceptible. This measure is widely employed in resistance screening studies. However, its downside is that it requires intensive processing of roots and, at times, soil, to recover specimens, usually carried out through centrifugal and sugar flotation as described by Coolen & D’Herde (1972). The difficulty of this method has led other phytonematode pathosystems to adopt alternative techniques, such as the gall index used to assess resistance to *Meloidogyne* spp., which induces easily visible root galls (Taylor & Sasser, 1978).

This difficulty in accessing RLN resistance is reflected in a scarcity of screening studies aimed at finding less susceptible genotypes across suitable crop hosts, with much of the literature on this matter coming from studies with other focuses, such as chemical treatments in which several varieties are tested, or smaller screening studies. This difficulty in assessing RLN resistance has resulted in a scarcity of dedicated screening studies to identify less susceptible genotypes among suitable crop hosts, particularly in sugarcane, where full resistance (RF ≤ 1) is not observed for *P.zeae*. Much of the available literature on this topic derives from studies with broader objectives, such as chemical treatment trials involving multiple varieties, or limited-scale screening efforts. From such studies, the two varieties examined here, CTC9001 and RB966982, were selected (Dinardo-Miranda et al., 2019; Santos et al., 2012). CTC9001 consistently exhibited higher susceptibility to *P. zeae* compared to other varieties in its screening panels, whereas RB966982 demonstrated a more resistant response relative to its counterparts in its respective evaluations. In light of these findings, CTC9001 and RB966982 are hereafter referred to as susceptible and resistant, respectively, in regard to *P. zeae* reproduction.

Therefore, based on the novel sugarcane genome assembly R570 (Healey et al., 2024), the aim of this study was to compare the transcriptional profiles of two sugarcane genotypes differing in their susceptibility to *Pratylenchus zeae* penetration, in order to uncover the molecular mechanisms underlying their responses to the pathogen. Our results reveal a set of genotype-specific mechanisms that may contribute to the differential success of nematode invasion.

## 3. MATERIALS AND METHODS

### 3.1 Selection and preparation of experiments for transcriptional profiling of sugarcane roots infected or not infected by *Pratylenchus zeae*

#### 3.1.1 Treatment of sugarcane setts

Based on a literature review, two contrasting *Saccharum* spp. varieties regarding susceptibility to *Pratylenchus zeae* infection were selected. Single-bud sugarcane setts from varieties RB966928 and CTC9001 were subjected to thermal and chemical treatments for decontamination (Fernandes Júnior et al., 2010). Initially, the setts were scrubbed using a sponge and then immersed in a water bath at 52°C for 30 minutes. After thermal treatment, the setts were transferred to a sodium hypochlorite solution composed of 950 mL of Milli-Q water and 50 mL of commercial bleach, where they remained for 10 minutes. Subsequently, the setts were rinsed three times with Milli-Q water to remove residual chemicals. The treated setts were placed in trays containing autoclaved vermiculite moistened with Milli-Q water and covered with perforated plastic to allow ventilation. The trays were transferred to a growth room and maintained at a constant temperature of 28°C, under a 12-hour light/12-hour dark photoperiod. The plastic covering was removed on the third day after the beginning of cultivation. During transplantation, the setts were rinsed with Milli-Q water to remove excess vermiculite and minimize contamination. The soil used was previously autoclaved and sieved to ensure sterility. Each sett, weighing an average of 30 grams (ranging from 10 to 15 grams), was transplanted into 300 mL disposable plastic cups filled with 150 grams of sterile soil. After transplantation, the soil was immediately moistened. The cups were placed in trays and transferred to the growth room at the Department of Genetics, ESALQ/USP, where they were maintained at 30°C under a 12-hour daily light cycle. Inoculation took place 24h after transplant. A total of a 1,000 *P.zeae* specimens of all life stages were evenly inoculated into two oblique holes of 2 and 4 cm flanking the transplanted sett in a 180° angle from each other.

#### 3.1.2 Sampling of infected and non-infected sugarcane plants

Sample collection was performed 15 days after transplantation, a timepoint established based on preliminary experiments in which consistent penetration was observed. Both roots and aboveground tissues were harvested. Roots were gently washed under running tap water using a sieve, followed by immersion in Milli-Q water to remove residual debris. Cleaned root tissues were placed into pre-labeled aluminum foil envelopes and immediately immersed in liquid nitrogen. Dissection of both roots and shoots was performed while the samples remained submerged in liquid nitrogen to prevent RNA degradation. The shoot tissue was defined as the stalks, excluding older leaves. All samples, grouped by biological replicate, were stored at –80°C until RNA extraction and further analysis.

#### 3.1.3 Root staining for nematode visualization

Collected roots were stained to visualize nematodes using acid fuchsin, following a modified protocol described by Byrd et al. (1983). Roots from the previously described treatments were selected for staining. Initially, roots were washed under running water using a sieve. They were then immersed for 5 minutes in a sodium hypochlorite solution (950 mL Milli-Q water + 50 mL commercial bleach). Afterward, roots were rinsed in Milli-Q water for 15 minutes to prepare for staining. Two staining reagents were used: acid fuchsin (pH 4.0) and acidified glycerol (pH 5.26). Roots were immersed in acid fuchsin inside a beaker and heated to boiling point. Once boiling was reached, roots were maintained in the solution for 30 seconds. They were then cooled to room temperature, washed with Milli-Q water, and transferred to a second beaker containing acidified glycerol. This solution was also heated to boiling point. Following this step, roots were drained and placed in Petri dishes to verify the presence of *P.zeae* lifestages under a light microscope at 15 DAI (Figure 1).

**Figure 1.**
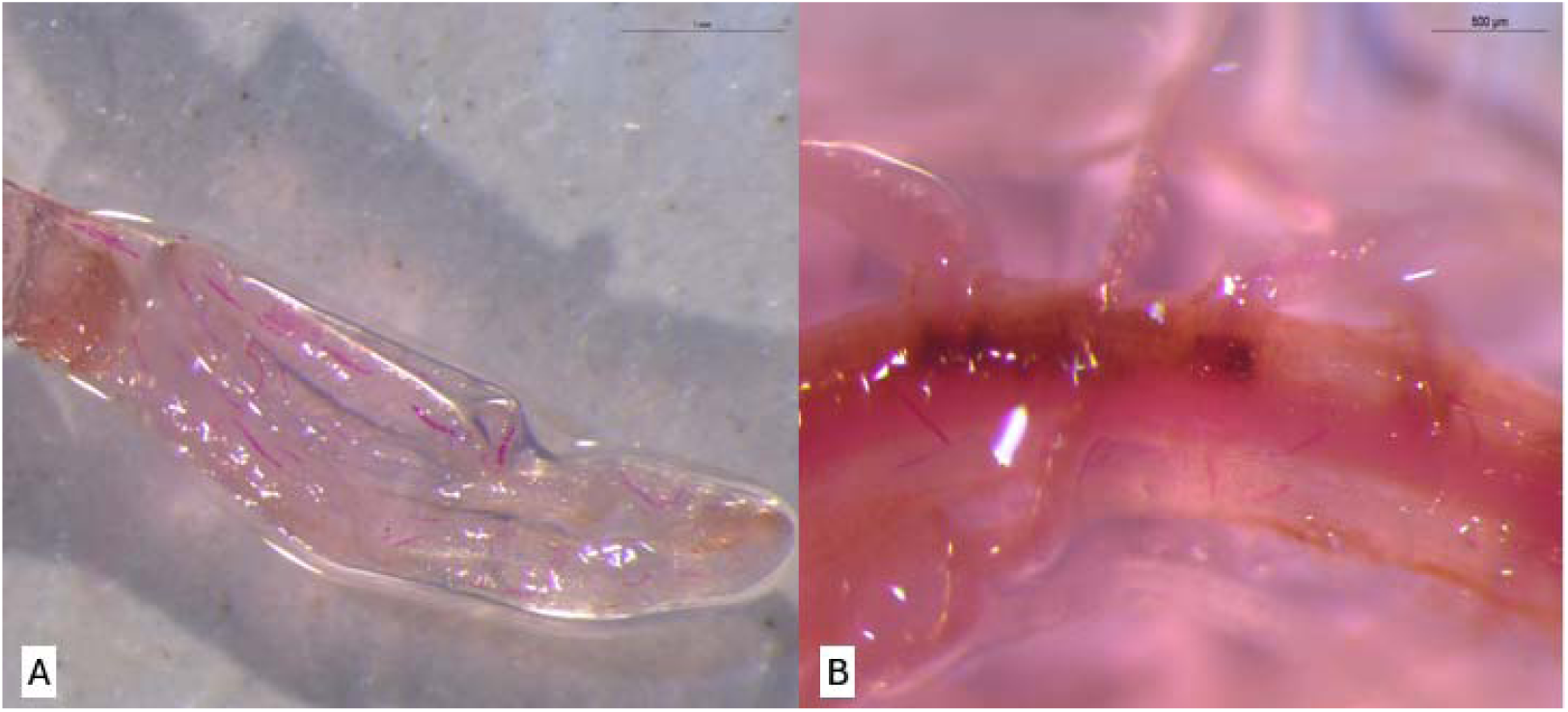
Visualization of *Pratylenchus zeae* life stages in sugarcane roots at 15 days after inoculation (DAI) using the acid fuchsin staining method described by Byrd et al. (1983). (A) Variety RB966928; (B) Variety CTC9001.

#### 3.1.4 Processing of collected root samples

Root maceration was performed using the CryoMill system (Retsch GmbH). The equipment was first pressurized to 1 atm to initiate the circulation of liquid nitrogen. Grinding jars and steel balls were pre-cooled to –196°C by immersion in liquid nitrogen. After each grinding cycle, the jars were washed with 70% ethanol and re-cooled before reusing to prevent cross-contamination. The CryoMill was programmed using memory slot P2 to enable nitrogen circulation prior to sample loading. Maceration was conducted using program P5, which applies a two-step grinding cycle: the first step circulates liquid nitrogen for 30 seconds at a frequency of 5 horizontal oscillations per second, followed by 30 seconds at a frequency of 30 oscillations per second. Fifty-milliliter grinding jars were used, and each biological replicate underwent three grinding cycles to ensure thorough homogenization. Throughout the procedure, the samples were maintained under constant exposure to liquid nitrogen to preserve RNA integrity. Following maceration, the powdered root tissue was transferred to labeled 50 mL Falcon tubes and stored at –80°C for downstream RNA extraction.

### 3.2 RNA extraction and DNAse treatment

Total RNA extraction for each biological replicate was performed using the PureLink RNA Mini Kit with TRIzol reagent (Invitrogen/Thermo, Cat. No. 12183018A), following the manufacturer’s instructions with minor adaptations. Prior to extraction, the centrifuge was pre-chilled to 4 °C. Two sets of 1.5 mL RNase-free microcentrifuge tubes and one set of 200 μL tubes were prepared. All reagents, including DEPC-treated 70% ethanol, were handled under RNase-free conditions. Root tissues (50–100 mg) were ground in liquid nitrogen and homogenized in 1 mL of TRIzol reagent. The lysates were incubated at room temperature for 5 minutes. Subsequently, chloroform was added, and the mixture was shaken manually for approximately 15 seconds, followed by incubation at room temperature for 3 minutes. Samples were centrifuged at 12,000 rpm for 15 minutes at 4 °C to separate phases. Approximately 600 μL of the upper, clear aqueous phase—containing RNA—was carefully transferred to a clean RNase-free tube. An equal volume of 70% ethanol was added to the aqueous phase, and the mixture was vortexed and inverted to avoid precipitation. Up to 700 μL of the mixture was loaded onto a spin column with a collection tube and centrifuged; this step was repeated until the entire sample passed through the column. The column was then washed with 700 μL of Wash Buffer I and centrifuged. This was followed by a second wash using 500 μL of Wash Buffer II. After a final centrifugation to dry the column, RNA was eluted by applying 30–100 μL of RNase-free water directly to the membrane. After a 1-minute incubation at room temperature, the column was centrifuged at 12,000 × g for 2 minutes. Eluted RNA was immediately stored at –80 °C. DNase treatment of the purified RNA was performed using the DNase I Amplification Grade Kit (Sigma-Aldrich). In a 200 μL tube, the total eluted RNA was treated with 1 μL of 10× Reaction Buffer, 1 μL of DNase I (1 U/μL), and up to 8 μL of RNA (leaving at least 2 μL reserved for quality control). The mixture was gently mixed and incubated at room temperature for 15 minutes. After incubation, 1 μL of Stop Solution was added, and the samples were incubated at 70[°C for 10 minutes in a thermal block to inactivate the DNase enzyme. Immediately afterward, tubes were transferred to ice for at least 5 minutes to ensure complete inactivation. Subsequent steps included RNA quantification, integrity assessment via gel electrophoresis, and storage at –80 °C for downstream analyses.

### 3.3 Sequencing and Transcriptome data processing

Library preparation was conducted according to manufacturer’s instructions of the Illumina® Stranded mRNA Prep Ligation kit, and RNA paired-end sequencing was performed by NextSeq 2000 (Illumina) in 2×100 bp runs at the Functional Genomics Center, ESALQ, University of São Paulo, Piracicaba, BR. Quality assessment of the raw RNA-seq reads (2×100 bp paired-end) was carried out using FastQC (Andrews, 2010) (v0.12.1) and processed with Cutadapt (v5.0) (Martin, 2011) to remove adapters and low-quality sequences. Cleaned reads were aligned to the *Saccharum spontaneum* × *S. officinarum* R570 reference genome (Healey et al., 2024) using HISAT2 (Zhang et al., 2021), and gene-level read counts were quantified with featureCounts (Liao et al., 2014).

### 3.4 Differential expression and functional enrichment analysis

Differentially expressed genes (DEGs) were identified using DESeq2 (v1.44.0) with a significance threshold of adjusted p-value (padj) < 0.05 and an absolute log2 fold change > 1 (Love et al., 2014). Four primary comparisons were carried out: (1) control vs. inoculated samples within the CTC9001 variety, (2) control vs. inoculated samples within the RB966982 genotype, (3) inoculated CTC9001 vs. inoculated RB966982, and (4) control CTC9001 vs. control RB966982. Prior to analysis, low-count genes (row sums < 10) were filtered out, and data were normalized to account for differences in library size. To assess overall sample relationships, we conducted principal component analysis (PCA) on rlog-transformed counts, incorporating 95% confidence ellipses to visualize biological replicates.

For functional interpretation, Gene Ontology (GO) enrichment analysis was performed using clusterProfiler (v4.12.0) (Yu et al., 2012) with the aid of GO.db (v3.19.1) (Carlson, 2017) database for term mapping. Enriched terms (false discovery rate, FDR < 0.05) were categorized in (BP/MF/CC) term rankings and data visualization was carried out by plotting gene-concept networks to highlight biological trends. All analyses were implemented in R (v4.3.1) (R a Language and Environment for Statistical Computing, 2010).

## 4. RESULTS

### 4.1 Transcriptional profiling of two sugarcane varieties in response to *P. zeae* infection

#### 4.1.1 Transcriptome sequencing and library assembly

Sequencing of the 12 paired-end samples yielded 389.8 million reads after adapter removal and ribosomal RNA depletion. Alignment rates to the R570 reference genome varied by treatment, depending on both variety and inoculation status. The CTC9001 variety exhibited lower mean alignment rates (55.6% and 65.1%), whereas RB966928 showed higher rates (88.0% and 84.9% for the control and inoculated treatments, respectively; Figure 2).

**Figure 2.**
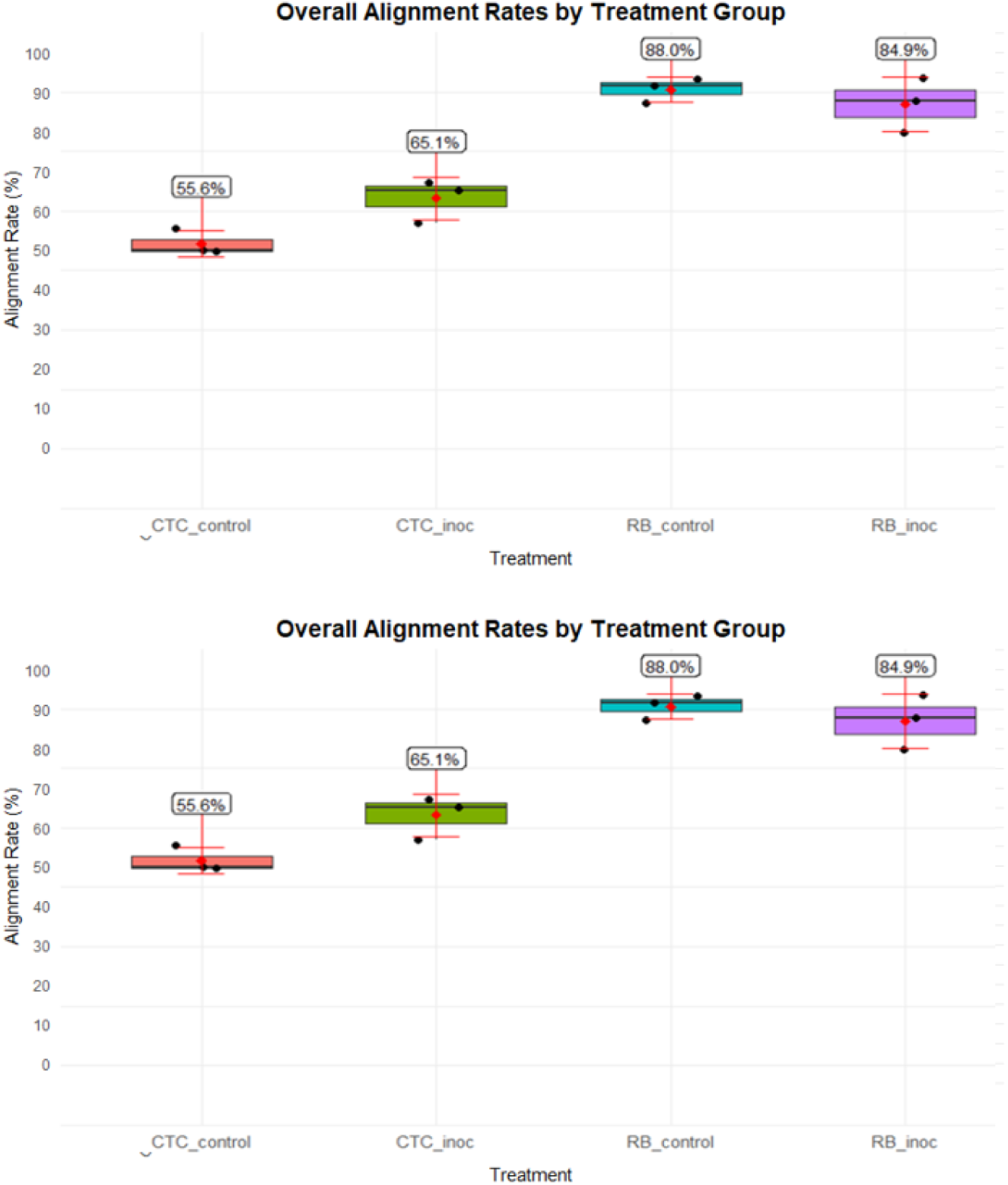
HISAT2 alignment percentages by experimental condition. Individual replicate data are indicated as black dots, while red points and bars illustrate means ±1 standard deviation (sd). Conditions shown include CTC_control, CTC_inoc, RB_control, and RB_inoc.

#### 4.1.2. Differential gene expression analysis of experimental contrasts

Principal component analysis (PCA) revealed a clear role of the sugarcane variety in PC1 (60% of the variance), while PC2 (26% of the variance) reflects the inoculation effect, with both inoculated treatments diverging from their respective control in the PC2, especially the resistant variety RB966982, which showed a strong transcriptional profile change upon inoculation (Figure 3).

**Figure 3.**
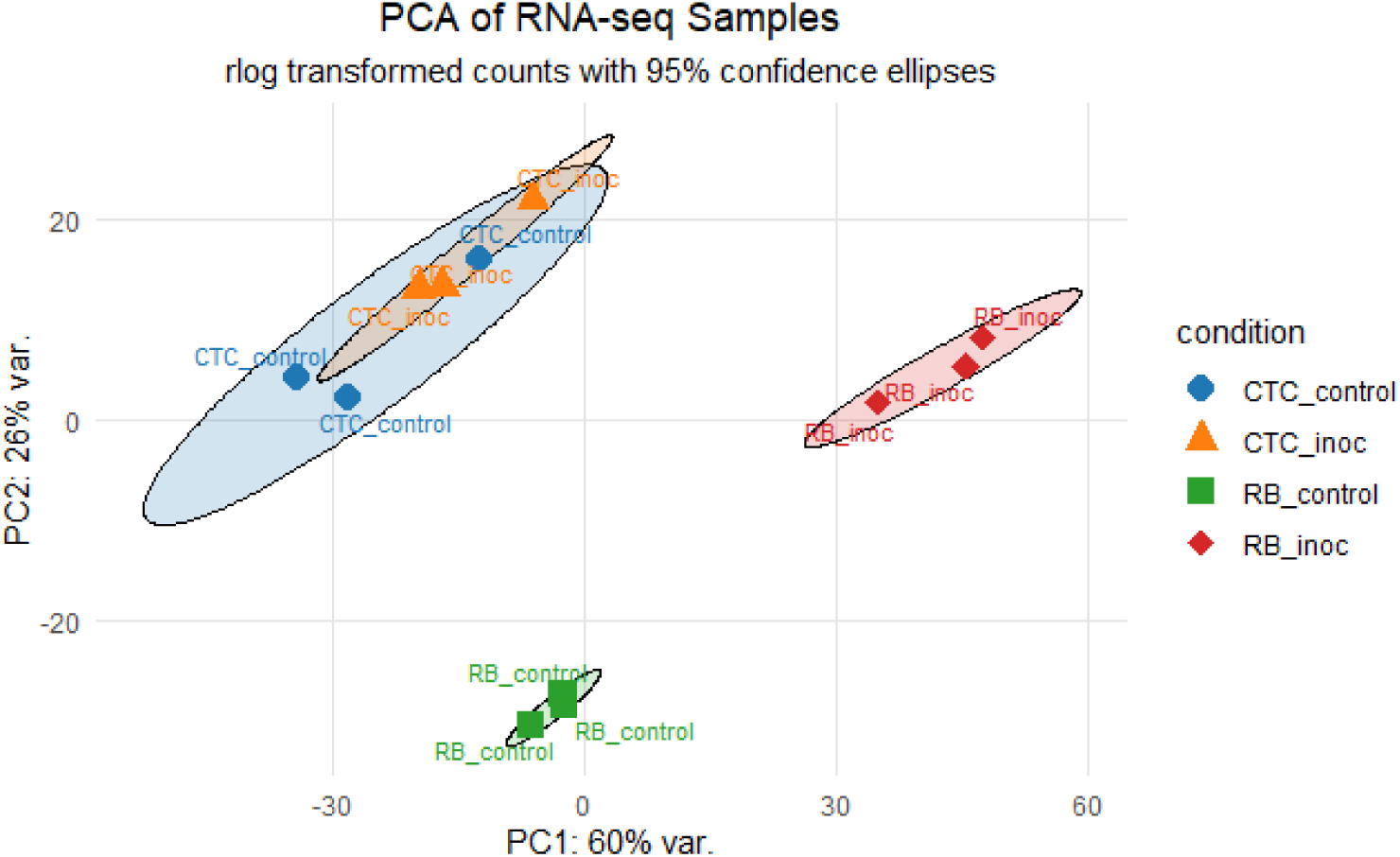
Principal component analysis (PCA) plot based on rlog-transformed RNA-seq count data, illustrating clustering patterns across samples. Ellipses indicate 95% confidence intervals for each group.

Principal component analysis (PCA) revealed a clear role of the sugarcane variety in PC1 (60% of the variance), while PC2 (26% of the variance) reflects the inoculation effect, with both inoculated treatments diverging from their respective control in the PC2, especially the resistant variety RB966982, which showed a strong transcriptional profile change upon inoculation (Figure 3).

Regarding DEGs, the CTC Control vs Inoculated contrast presented a modest transcriptional reprogramming, with 1,426 upregulated and 1,959 downregulated genes, in strong contrast to the RB Control vs Inoculated treatment, which displayed 5,334 upregulated and 3,355 downregulated genes. Additionally, in the analysis of inter-variety comparisons, CTC vs RB under control conditions revealed 10,859 upregulated and 9,320 downregulated transcripts, indicating broad constitutive differences between the two genotypes. The strongest transcriptional divergence was observed under inoculated conditions, with 16,534 upregulated and 14,819 downregulated genes, highlighting that infection further accentuates genotype-specific differences (Figure 4).

**Figure 4:**
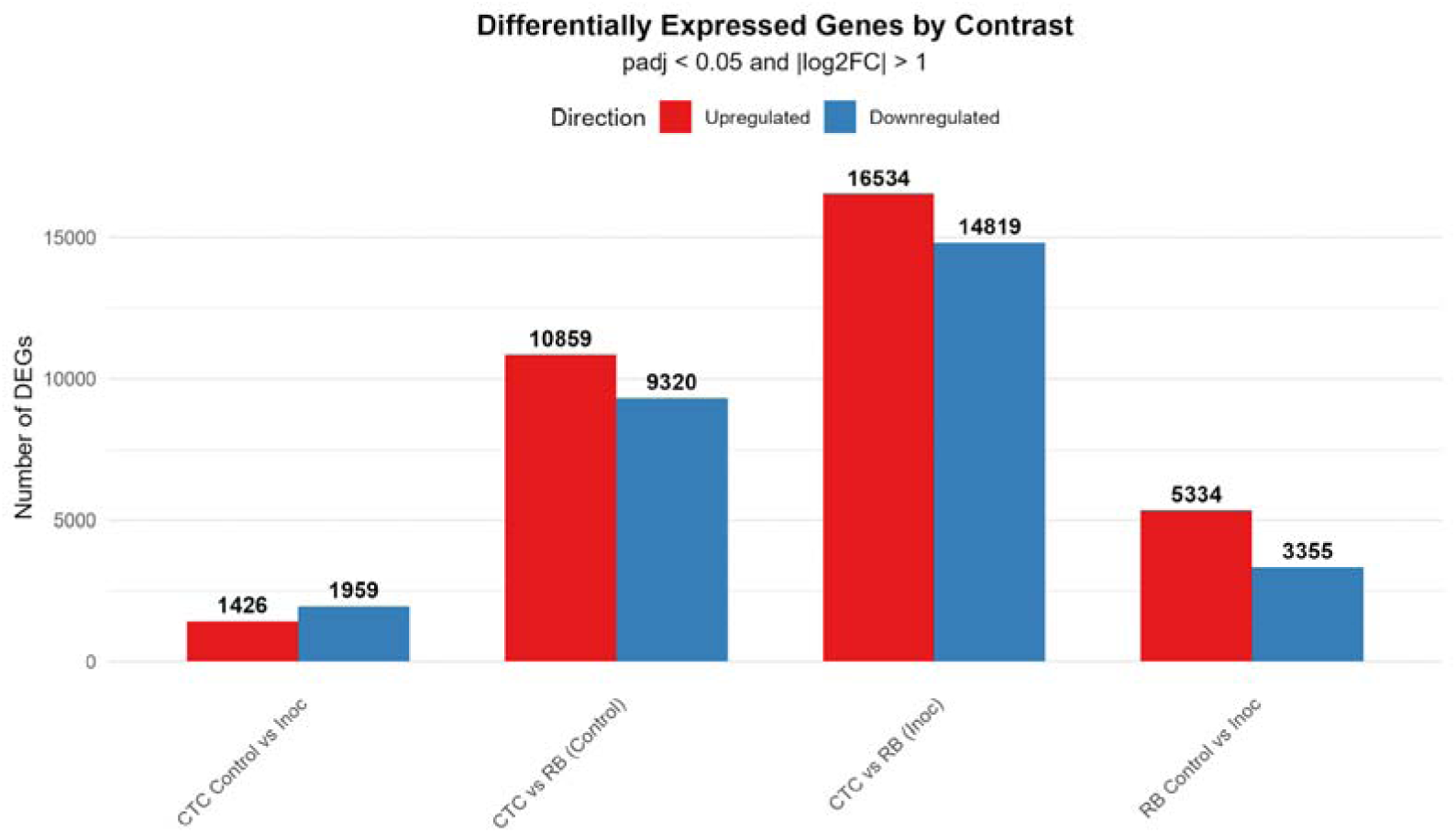
Bar plot displaying the number of significantly upregulated (red) and downregulated (blue) genes across four pairwise comparisons. Differentially expressed genes (DEGs) were identified using a threshold of adjusted *p*-value (padj) ≤ 0.05 and an absolute log_₂_ fold-change of (|log_₂_FC|) > 1.

#### 4.1.3. Gene ontology enrichment analysis and co-expression networks

Enriched GO terms (FDR < 0.05) were found across all contrasts, including the inter-variety control comparison CTC_control x RB_control (mock-inoculated) at 15 DAI. Overall, molecular function (MF) terms were prevalent across all contrasts, followed by biological processes (BP) and cellular component (CC). The varying number of total unique enriched GO terms in each contrast highlights a substantially distinct within-variety response upon nematode infection. For instance, the CTC_inoc vs RB_inoc contrast showed the most robust response with 314 unique GO terms, followed by the RB_control with 272 terms. It is noteworthy that despite presenting the second highest number of DEGs (20 179), the CTC_control vs RB_control has the lowest number of unique enriched GO terms (148), which is closely matched only by the CTC_control vs CTC_inoc with 154 GO terms, this one however, presented the least amount of DEG’s (3 385). These inversely proportional patterns highlight a strong transcriptional bias toward specific metabolic shifts upon inoculation across all contrasts when the nematode is present.

Data visualization through Concept Network Plot (cnetplot) of the top 8 ranked GO terms for each contrast and direction (induced/up or repressed/down), showed significantly distinct response patterns upon nematode infection depending on the variety, and also confirmed the underlying discrepant basal expression pattern between each variety’s control presented in the PCA and DEGs analysis.

##### 4.1.3.1 Intra-variety analysis

This section presents the results of GO terms enriched upon *P. zeae* inoculation at 15 DAI. Enriched networks were classified as induced or repressed based on the up-or downregulation patterns of their gene components, using the respective mock-inoculated controls as the reference.

###### Susceptible interaction (CTC_control vs CTC_inoc)

The up-regulation network consisted mainly of three prominent hubs as seen in Figure 5, the densest one interconnected groups of genes with microtubule-based movement annotations (GO:0007018, BP), all anchored by kinesin complex (GO:0005871, CC), microtubule motor activity (GO:0003777, MF) and microtubule binding activity (GO:0008017, MF). The second most overrepresented hub consisted of cell wall polysaccharide synthesis elements, composed of a cluster of genes involved in 1,3-β-D-glucan synthase activity (GO:0003843, MF) together with the 1,3-β-D-glucan synthase complex (GO:0017011, CC) and the (1→3)-β-D-glucan biosynthetic process (GO:0051274, BP). A lone hub centered on cytoskeletal motor activity (GO:0003774, MF) can also be observed. The down-regulation network for this contrast is dominated by a hub consisting of nutrient reservoir activity (GO:0045735, MF) and manganese ion binding (GO:0030145, MF), also present here is a hub of down-regulated defense response genes in clusters corresponding to defense response to bacterium (GO:0042742, BP) and defense response to fungus (GO:0050832, BP). Non-connected hubs here are represented by a module comprising the eukaryotic translation elongation factor 1 complex (GO:0005853, CC) along with asparagine synthase (glutamine-hydrolyzing) activity (GO:0004066, MF), underscoring reduced protein synthesis and amino-acid metabolism (Figure IV).

**Figure 5.**
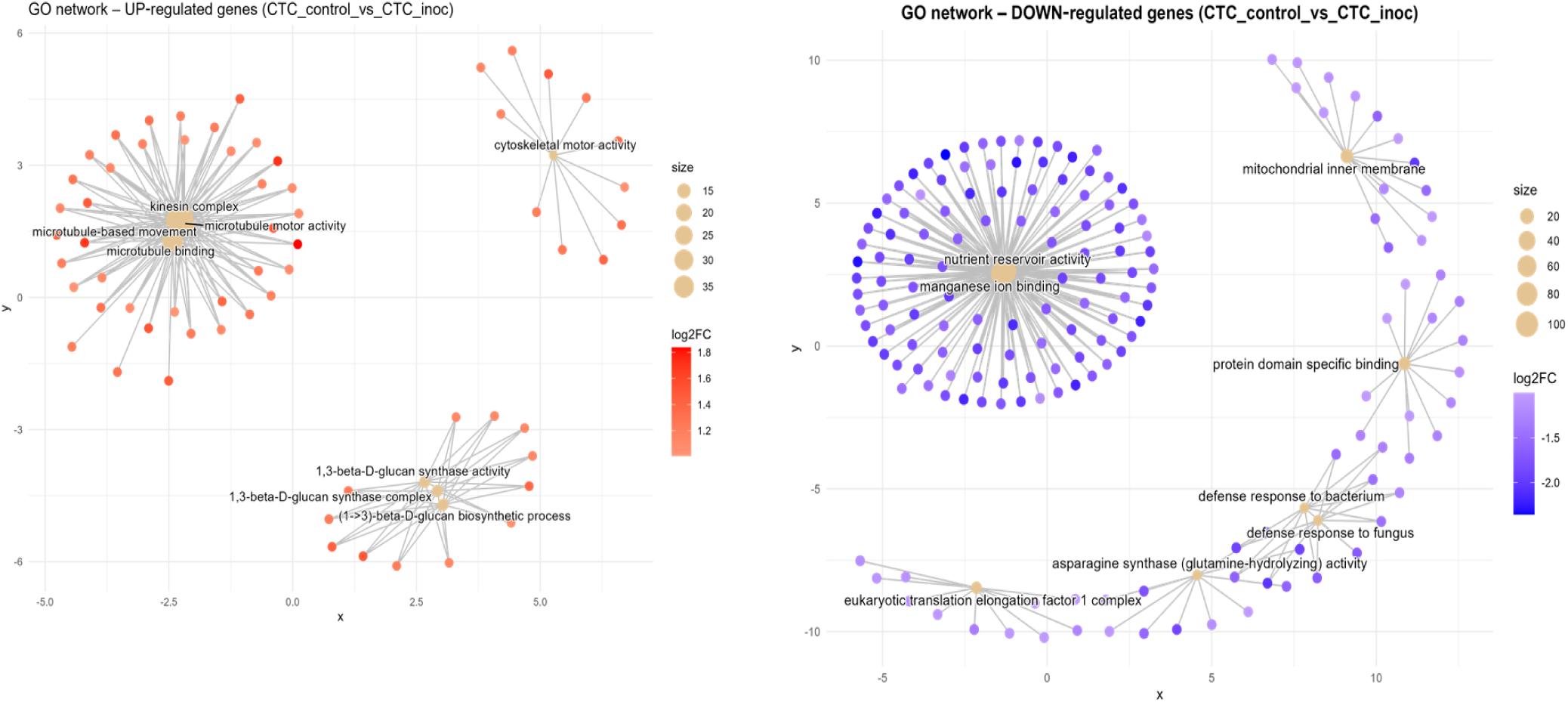
Concept network plots illustrating GO term enrichment for significantly upregulated (left, red nodes) and downregulated (right, blue nodes) genes upon inoculation in the susceptible variety CTC9001 at 15 DAI. Node size represents the number of associated genes within each GO term, and node color intensity corresponds to the magnitude of differential expression (log2 fold-change). Lines represent the relationships between genes and their respective enriched GO terms.

###### Resistant interaction (RB_control vs RB_inoc)

In the resistance response to nematode infection, induced networks of GO terms formed several hubs (Figure 6). First, a complex central network containing an abundance of genes related to glycolytic process (GO:0006096, BP) can be observed. In one adjacency of this term is a tightly connected cluster for 6-phosphofructokinase activity (GO:0003872, MF) and the fructose-6-phosphate metabolic process (GO:0006000, BP), all related to early steps in sugar catabolism. On the opposite side, the glycolytic process group is connected to a broad grouping around oxidoreductase activity, acting on single donors with incorporation of molecular oxygen (GO:0016701, MF). To the right, genes linked to malate dehydrogenase (decarboxylating) (NAD) activity (GO:0004470, MF) form a discrete module linked closely with a cluster for malic enzyme activity (GO:0004477, MF). To the bottom left two gene hubs are connected under the enriched GO terms thiamine pyrophosphate binding (GO:0031993, MF) and carboxylyase activity (GO:0016831, MF), signaling decarboxylation/carboxylation metabolism. A lone hub consisting of galactoside 2-alpha-L-fucosyltransferase activity (GO:0015031, MF) is also present, indicating cell-wall modification activity (Figure 6). Conversely, down-regulated networks reveal a suppression of photosynthetic processes in the resistance response, with a sprawling module for photosynthesis (GO:0015979, BP) interconnecting with photosystem I reaction center (GO:0009522, CC) and the thylakoid membrane (GO:0042651, CC). Additionally, a lone neighboring hub for photosynthesis, light harvesting (GO:0009765, BP) is present here. O-methyltransferase activity (GO:0008171, MF) forms its own cluster, while a group for serine-type endopeptidase inhibitor activity (GO:0004867, MF) is formed together with response to wounding (GO:0009611, BP). A two-component hub consisting of genes annotated to amino acid metabolic process (GO:0006520, BP) and carboxylase activity (GO:0016831, MF) are also present here.

**Figure 6.**
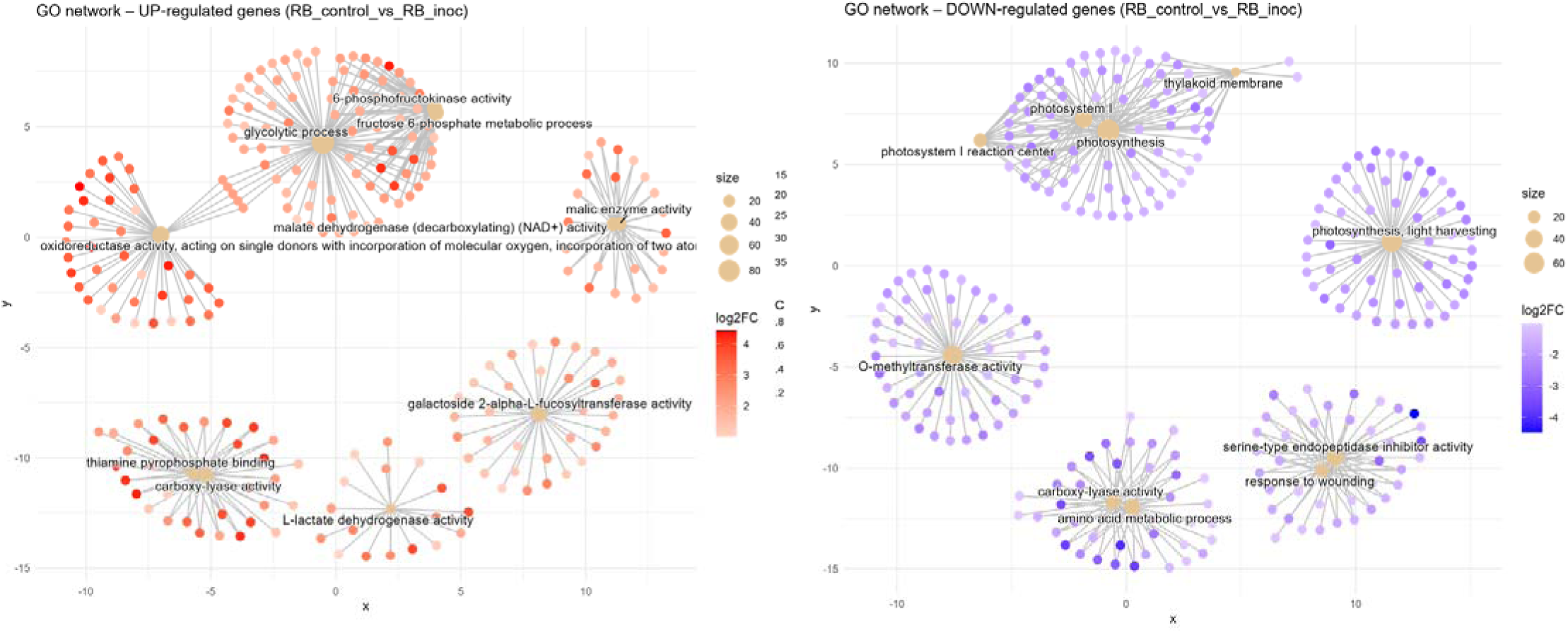
Concept network plots illustrating GO term enrichment for significantly upregulated (left, red nodes) and downregulated (right, blue nodes) genes upon inoculation in the resistant variety RB966982 at 15 DAI. Node size represents the number of associated genes within each GO term, and node color intensity corresponds to the magnitude of differential expression (log2 fold-change). Lines represent the relationships between genes and their respective enriched GO terms.

##### 4.1.3.2 Inter-variety analysis

Here the two inter-variety contrasts are compared, first in response to the pathogen inoculation (CTC_inoc x RB_inoc), and second in response to the mock inoculation (CTC_control x RB_control), to compare overall expression profiles in the absence of the biotic stress caused by the nematode. As there are no controls, the comparisons made here are based on comparative transcriptional levels, and results are presented as induced in comparison to each other. This dual approach was conducted to observe both stress-responsive and baseline transcriptional differences between varieties.

###### Susceptible vs resistant direct comparison upon inoculation (CTC_inoc x RB_inoc)

When compared to its susceptible counterpart, the RB variety induced terms network coalesced into five major functional hubs (Figure 7). First, a central module consisting glycolytic process (GO:0006096, BP) was tightly connected to an oxidoreductase activity, acting on single donors with incorporation of molecular oxygen (GO:0016701, MF) cluster, indicating a coordinated up-regulation of sugar catabolic and redox enzymes. In its immediate adjacency, a hub containing genes annotated to the thiamine pyrophosphate binding term (GO:0031993, MF) highlights the activation of thiamine dependent processes. Positioned at the lower left, a second network of genes centered on fatty acid biosynthetic process can be observed (GO:0006633, BP), reflecting upregulation of lipid metabolism. In the bottom right, two connected networks can be observed sharing similarly annotated genes related to peptide-methionine (R)-S-oxide reductase activity (GO:0050626, MF) and cellular response to stimulus (GO:0051716, BP), indicating an elevated protein repair and stress sensing machinery, close to these, a lone hub representing cysteine-type endopeptidase inhibitor activity (GO:0004869, MF) can be observed.

**Figure 7.**
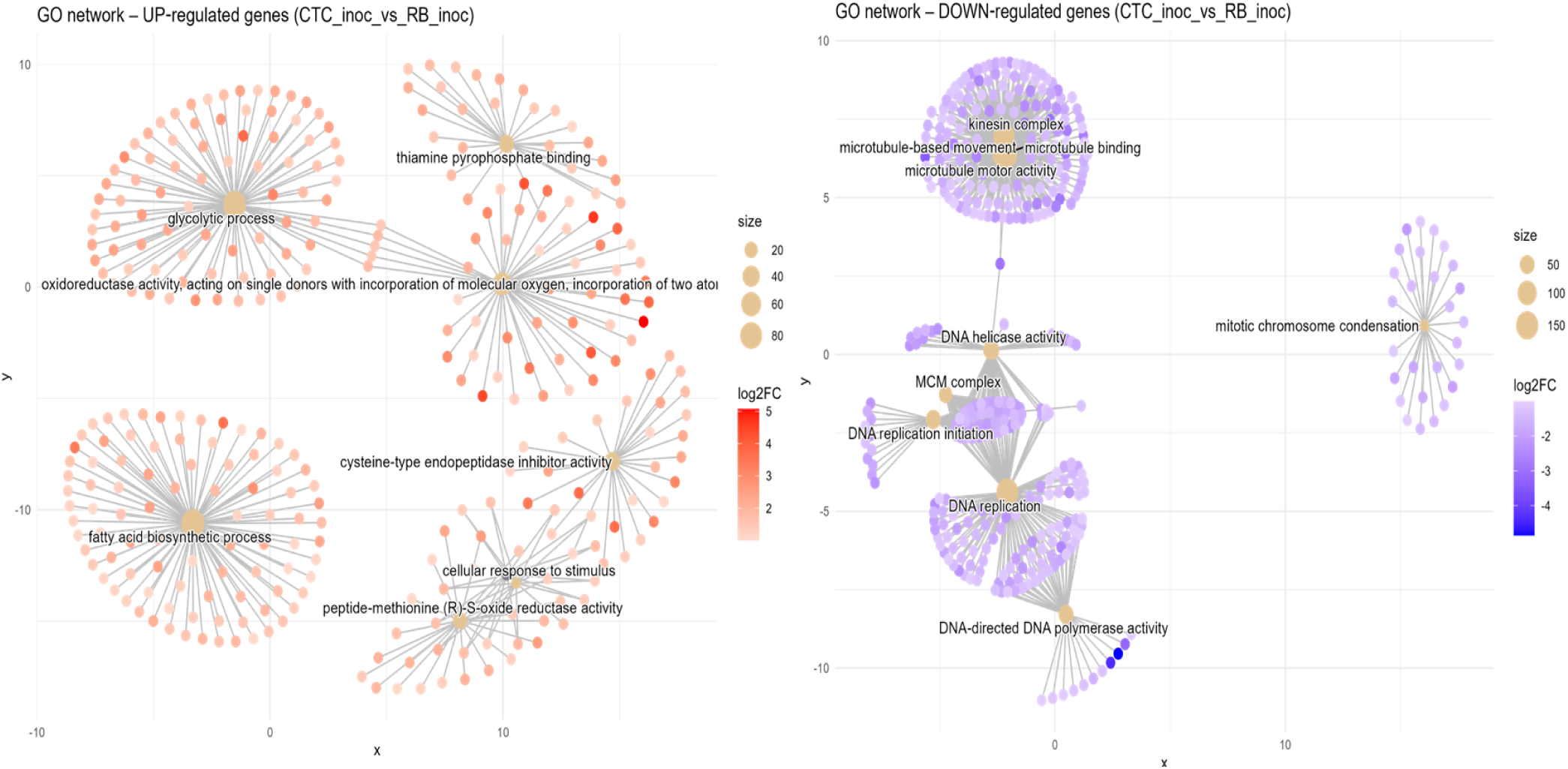
Concept network plots illustrating GO term enrichment for genes significantly induced in resistant variety RB966982 (left, red nodes) and comparatively induced in susceptible variety CTC9001 (right, blue nodes), upon inoculation at 15 DAI. Node size represents the number of associated genes within each GO term, and node color intensity corresponds to the magnitude of differential expression (log2 fold-change). Lines represent the relationships between genes and their respective enriched GO terms.

In the susceptible response, the CTC variety inverse direction of enriched go terms presents networks of induced genes divided into several GO terms spread across four discrete hubs. The top left module is centered on microtubule-based movement (GO:0007018, BP) and is linked to the terms: microtubule motor activity (GO:0003777, MF), kinesin complex (GO:0005871, CC) and microtubule binding (GO:0008017, MF), which indicates a coordinated activation of cytoskeletal transport machinery. The hub below is anchored by DNA helicase activity (GO:0003678, MF) which connects to the MCM complex (GO:0042555, CC) and DNA replication initiation (GO:0006270, BP), reflecting up-regulation of replisome assembly factors. The bottommost hub is centered around a dense cluster of DNA replication GO annotated genes (GO:0006260, BP), closely related to DNA-directed DNA polymerase activity (GO:0003968, MF), highlighting enhanced genome duplication. A lone module is defined by mitotic chromosome condensation (GO:0007076, BP), which indicates mitosis related processes. Overall, the networks here represented marks GO terms closely associated to cellular multiplication in the susceptible response.

###### Susceptible vs resistant direct comparison of mock-inoculated expression (CTC_control x RB_control)

In the absence of the pathogen, the RB mock inoculated variety presents five distinct functional hubs that are induced in contrast with the CTC mock inoculated variety (Figure 8). First, a central module is tied together by xyloglucan metabolic process (GO:0010411, BP) which is tightly interconnected with xyloglucan:xyloglucosyl transferase activity (GO:0016762, MF) and apoplast (GO:0048046, CC), pointing to cell wall altering activities. In the rightmost upper corner is a hub organized around genes associated to lipid transport (GO:0006869, BP) highlights active metabolism of fatty acids and/or sterols, which might support membrane repair or signal transduction. To the bottom right the term oxidoreductase activity, acting on single donors with incorporation of molecular oxygen (GO:0016701, MF), highlighting the recurrence of this GO term across RB contrasts, even in the absence of *P. zeae*, this term captures up-regulation of ROS-generating and detoxifying enzymes. Two more enriched terms are found at the bottom left, one is DNA-3-methyladenine glycosylase activity (GO:0003884, MF) and the other is, phospholipase A activity (GO:0032435, MF), pointing to a basal tendency to DNA repair activity and lipid remodeling/repurposing metabolism, respectively.

**Figure 8.**
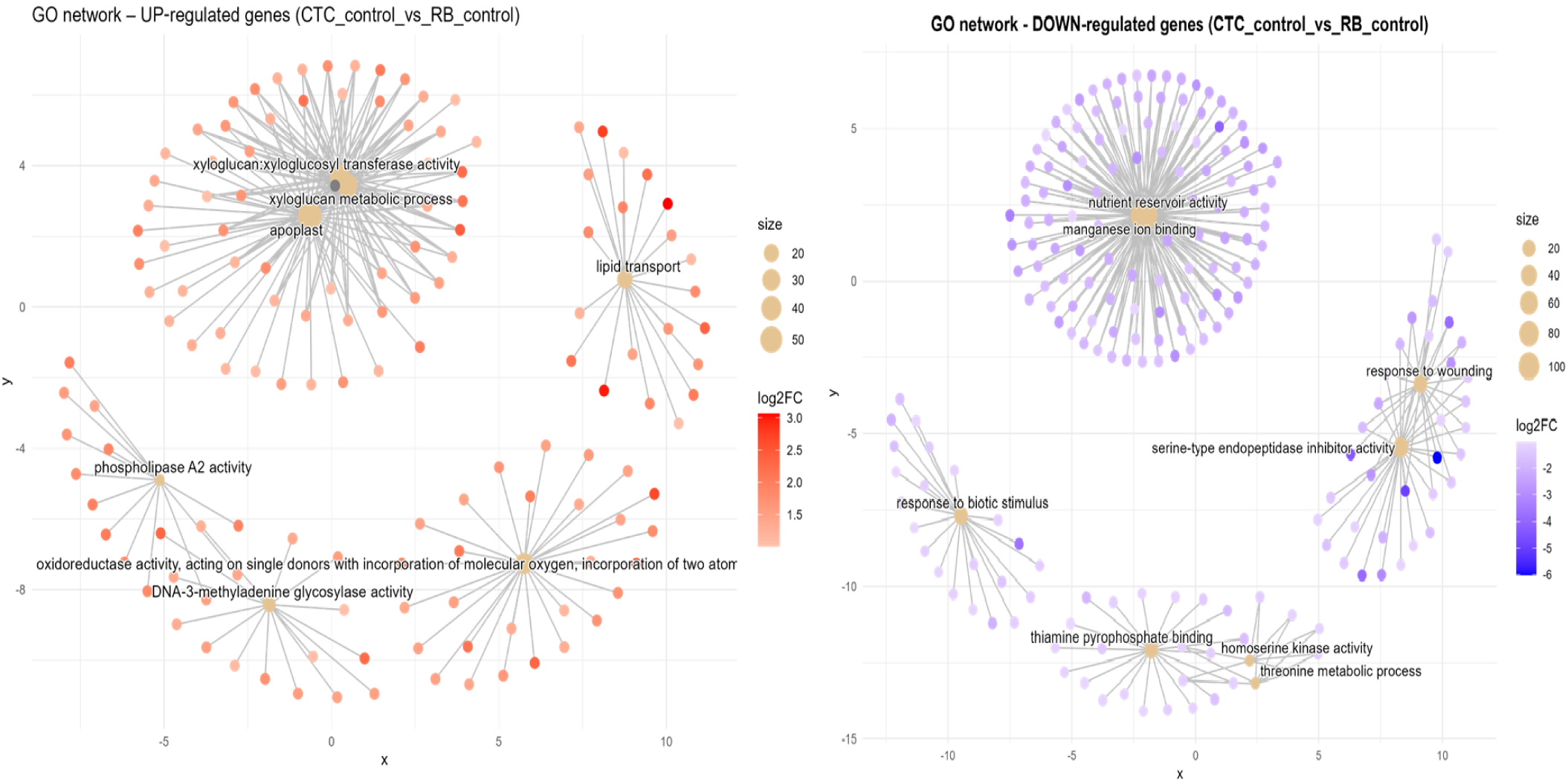
Concept network plots illustrating GO term enrichment for genes significantly induced in resistant variety RB966982 (left, red nodes) and comparatively induced in susceptible variety CTC9001 (right, blue nodes), upon inoculation at 15 DAI. Node size represents the number of associated genes within each GO term, and node color intensity corresponds to the magnitude of differential expression (log2 fold-change). Lines represent the relationships between genes and their respective enriched GO terms.

Compared to its mock inoculated RB counterpart, the CTC_control presents a central network hub consisting of nutrient reservoir activity (GO:0045735, MF) together with manganese ion binding (GO:0031402, MF), this hub was mentioned before in the CTC_control x CTC_inoc contrast as being repressed, here they are induced, which highlights a natural tendency to accumulate reserves in the CTC variety, which is repressed upon nematode infection (Figure 8). Interestingly, a cluster of defense related terms is formed by two networks, one, containing two hubs consists of response to wounding (GO:0009611, BP) and serine-type endopeptidase inhibitor activity (GO:0004867, MF), the other is a discrete network composed of genes related to response to biotic stimulus (GO:0009607, BP). Additional enriched terms here are present at the bottom center, they are thiamine pyrophosphate binding (GO:0019255, MF), homoserine kinase activity (GO:0004415, MF) and threonine metabolic process (GO:0006564, BP).

#### 4.1.4. Heatmap of normalized expression of resistance gene analogs reveals distinct basal and infection-induced defense patterns

To observe patterns in the expression of resistance gene analogs (RGA) across all contrasts, normalized counts for NB-ARC and NBS-LRR type genes were hierarchically clustered into two main modules (Figure 9). From top to bottom, the first module represents genes with low basal expression in CTC9001 that are slightly activated upon nematode inoculation; conversely, these same genes show higher basal expression in RB966982 and undergo robust up-regulation upon inoculation. The second module comprises two distinct clusters: the upper cluster consists of several genes with higher basal expression in CTC9001, most of which are further induced by nematode presence, while these loci remain repressed in RB966982 under both conditions. The lower cluster contains genes expressed at similar levels in CTC9001 regardless of treatment, but with basal repression in RB966982 and significant induction only upon nematode inoculation. Together, these patterns suggest that RB966982’s resistance is based on both a constitutive reservoir of expressed R-genes (upper module) and a selective induction of others (lower secondary cluster), whereas CTC9001 presents a robust expression of RGAs that are seemingly less effective to mount a resistance against *P. zeae*.

**Figure 9.**
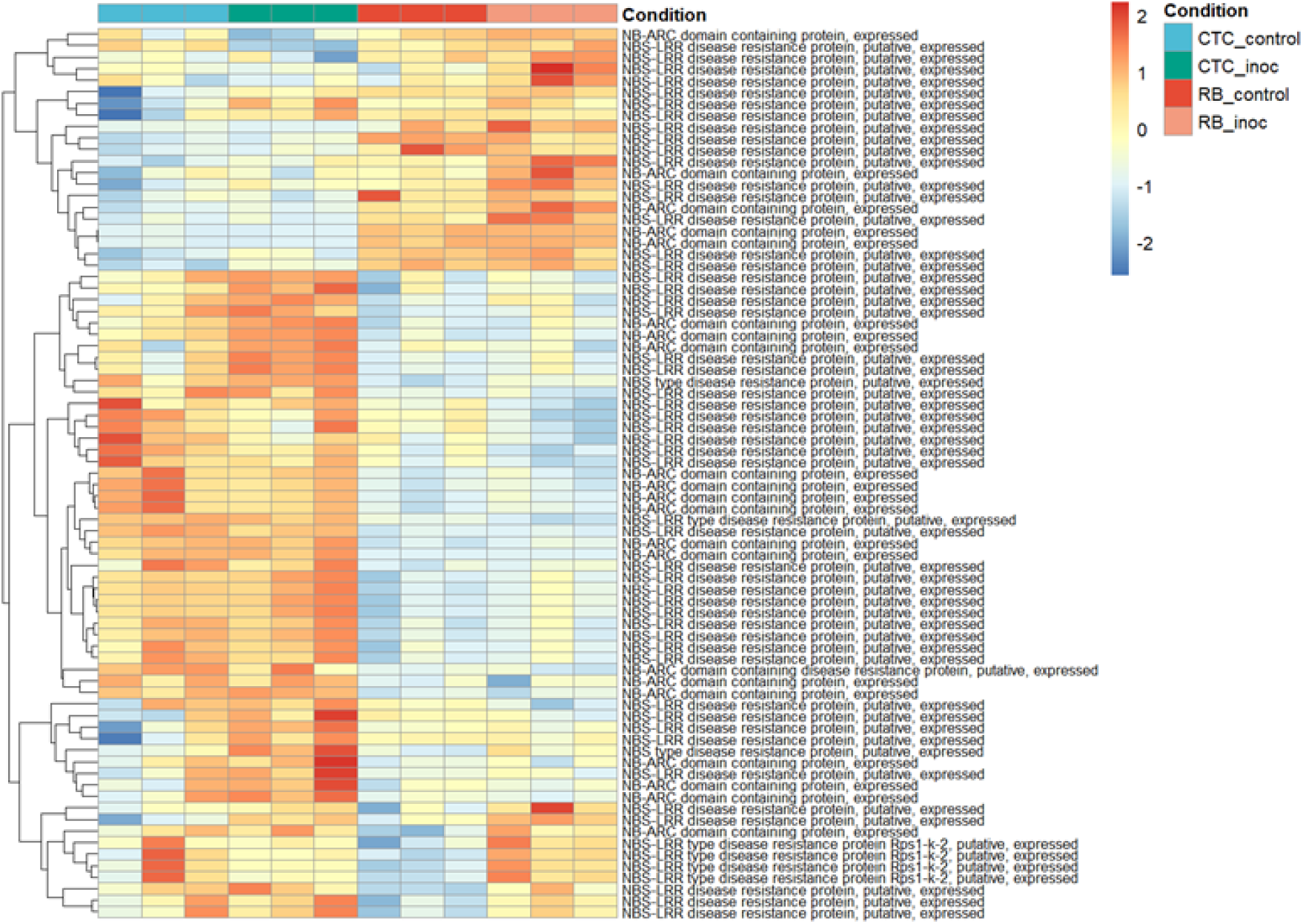
heatmap representing normalized expression (Z-score) of annotated genes involved in resistance responses across susceptible (CTC9001) and resistant (RB966982) varieties at 15 DAI. Color scale indicates relative expression levels ranging from highly induced (red) to strongly repressed (blue). Gene clustering (rows) reflects similarity in expression patterns across samples (columns).

Additionally, three RGAs were observed with interesting up-regulation patterns in the CTC_inoc × RB_inoc comparison, a contrast that allowed us to capture genes not transcribed (or barely transcribed) in one of the varieties. Specifically, SoffiXsp0nR570.05Dg100600, SoffiXsp0nR570.09Eg112400 and SoffiXsp0nR570.09s19063800 (Figure 10) displayed distinct behaviors: SoffiXsp0nR570.05Dg100600 (LFC:2.7) showed low basal expression in CTC9001 with a modest increase upon inoculation, yet in RB966982 its already higher basal levels climbed further after infection. In contrast, SoffiXsp0nR570.09Eg112400 (LFC:6.2) and SoffiXsp0nR570.09s19063800 (LFC:5.9) were virtually undetectable in CTC9001 but were constitutively expressed at moderate levels in RB966982 and then strongly induced by nematode challenge. These observations strengthen the perception that RB966982 combines a pre-existing reservoir of RGA transcripts with an infection-triggered amplification in a more successful manner than the CTC9001 variety in mounting a defense response against this nematode (Figure 10).

**Figure 10.**
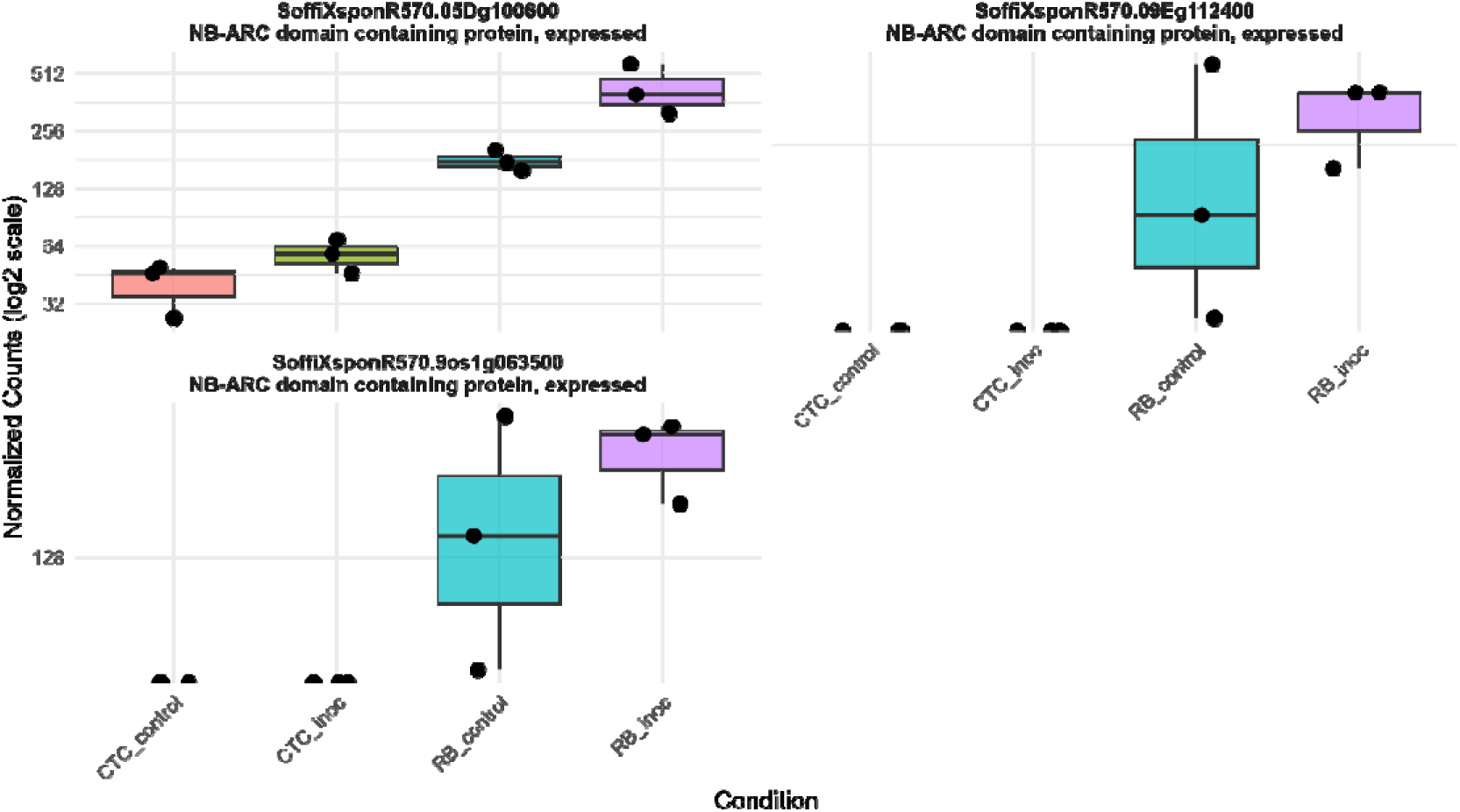
Boxplots illustrating expression patterns (normalized counts) of three distinct RGAs across all contrasts sugarcane varieties under control and inoculated conditions at 15 DAI.

#### 4.1.5. Divergent metabolic defense programs differentiate sugarcane varieties RB966982 and CTC9001 during nematode challenge

To summarize key defense strategies that will be presented in this section, Figure 11 highlights the transcriptional re-programming across six afore-mentioned GO-enriched pathways when the two sugar-cane varieties (CTC9001 and RB966982) are challenged with the pathogen. The panel to the left (intra-variety) shows how each genotype reacts to infection. In which CTC mounts a pronounced 1,3-β-D-glucan synthase response (12 genes induced, none repressed), whereas RB leaves this pathway essentially untouched and instead boosts xyloglucan α-L-fucosyltransferase (34 genes induced, one repressed). In other words, CTC reinforces callose deposition, while RB remodels xyloglucan branches—two alternative strategies for strengthening the cell wall.

**Figure 11.**
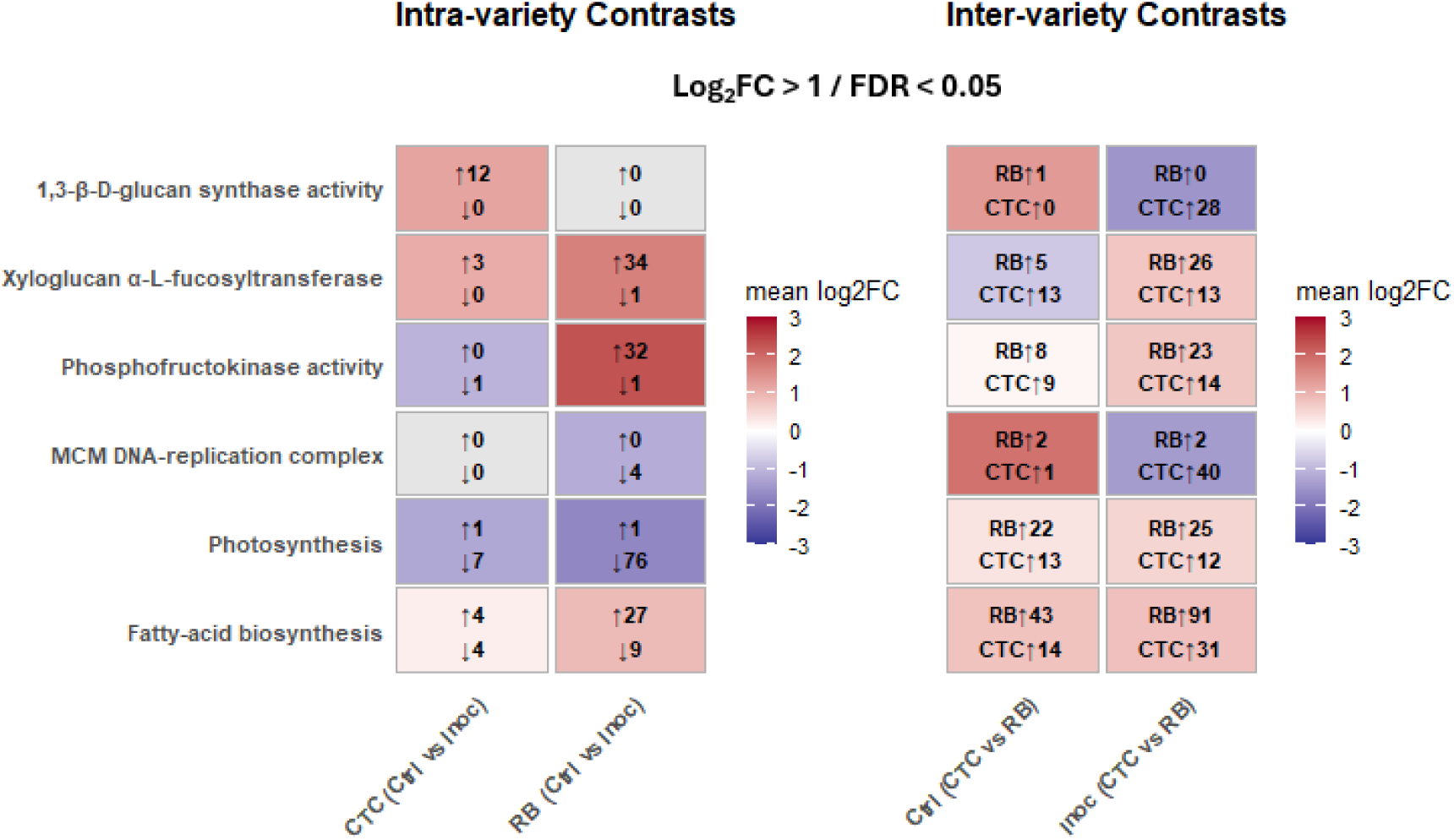
Overall of mean log_₂_ fold-changes (log_₂_FC > 1 / FDR < 0.05) for six GO-enriched pathways in intra-variety (CTC and RB, Control vs Inoculated) and inter-variety (Control and inoculated, RB vs CTC) contrasts. Within the intra-variety panel, red shading highlights gene sets with a positive mean log_₂_FC (up-regulated), while blue shading highlights those with a negative mean log_₂_FC (down-regulated). Arrows indicate the number of up-regulated and down-regulated genes meeting the cutoff within each set. In the inter-variety panel, red shading highlights pathways more induced in RB, blue in CTC; arrows indicate RB and CTC gene counts within each set.

For phosphofructokinase (PFK), an indicator of glycolytic intensification, RB exhibits a massive burst (32 genes up, one down), contrasting with CTC’s marginal change (one gene down). This indicates that RB diverts substantial carbon into glycolysis upon nematode infection.

Growth-related processes are regulated in opposite directions. RB strongly represses both the MCM DNA-replication complex (four genes down) and photosynthesis (76 genes down), prioritizing defence over proliferation and energy capture. CTC shows only mild repression of these pathways during infection, but in the inter-variety comparison the situation flips: many MCM and photosynthetic transcripts are still more abundant in RB than in CTC, hinting that RB entered the stress episode with a higher basal expression level. Finally, fatty-acid biosynthesis is moderately modulated in CTC (4 up, 4 down) but markedly induced in RB (27 up, 9 down intra-variety; 91 up versus 31 in CTC under infection).

Taken together, the heatmaps reveal two contrasting defence directives: CTC rapidly activates β-glucan synthesis and later compensates with DNA replication, whereas RB reallocates resources from growth and photosynthesis toward glycolysis, xyloglucan remodeling, and fatty-acid production.

#### 4.1.5. Distinct cell wall modification strategies are triggered in contrasting varieties upon inoculation with *P. zeae*

As reported in the GO enrichment section, distinct gene networks related to cell wall modifications were induced in each sugarcane variety upon nematode inoculation. While the CTC variety induced the transcription of genes involved in the 1,3-β-D-glucan synthesis upon nematode inoculation, the RB variety increased the number of transcripts involved in fucosyltransferase activity. To visualize this relationship, a heatmap was generated to display DEGs with annotated functions in both pathways across all experimental contrasts. Regarding 1,3-β-D-glucan synthesis, despite one of the replicates showing higher basal transcription rates for this gene group, the overall CTC_control group remains below the mean (blue tones). After inoculation, the CTC_inoc samples clearly shift towards higher Z-scores (approximately 1.0–1.5), as indicated by warmer colors (sup. Fig 1). This pronounced induction in CTC contrasts with the RB response, which exhibits only modest up-regulation in seven out of nine genes and down-regulation in two out of nine genes, with the majority of RB samples consistently remaining below a Z-score of zero.

Regarding xyloglucan fucosyltransferases, despite some basal variation among replicates, the overall CTC_control group shows predominantly negative Z-scores (blue tones), indicating below-average basal expression. Upon inoculation, CTC_inoc samples presents a modest shift toward neutral or slightly positive Z-scores (white to pale pink tones), suggesting a mild induction, especially in the cluster of genes at the bottom of the heatmap. This pattern contrasts sharply with the RB cultivar, which exhibits relatively low/neutral basal levels in RB_control and a very pronounced induction in RB_inoc, reflected by consistently high positive Z-scores (deep red). This robust activation is observed across most of the genes within this group, highlighting a cultivar-specific transcriptional response, potentially implicating xyloglucan fucosyltransferases as important players in the resistance mechanism observed in RB966982 (sup. Fig 2). Altogether, these data reveal genotype-specific strategies for either regulation of β-1,3-glucan deposition during the immune response or the activation of cell-wall fucosylation machinery in response to *P.zeae* infection.

#### 4.1.6. An intense shift in energy metabolism can be observed in the RB966982 variety 15 DAI

While the susceptible CTC variety presented a response focused mainly on the up-regulation of 1,3 beta-glucans, cellular transportation and cytoskeleton activity, the RB variety presented a more robust response. At the center of this response several genes connected to energetic metabolism can be observed, these are involved mainly in glycolysis and oxireductases up-regulation. (Figures 6 and 7). This shift is highlighted in the (sup. Fig 7), which provides a clear picture of how the glycolytic-process is profoundly altered based on a heatmap of its respective enriched GO term (GO:0006096). In synthesis, upon inoculation, CTC_inoc replicates shift into neutral or mildly positive (Z-scores of 0 to 0,5) for a small group of genes, indicating a modest activation of glycolysis in response to *P. zeae* infection. Conversely, the resistant RB966982 displays low basal levels of these same enzymes in RB_control (Z-scores < 0), and RB_inoc samples push almost every gene in the pathway into strongly positive z scores over 1 (bright reds). This consistent pattern across key genes including 6-phosphofructokinases, pyruvate kinases, enolases, among others, reveals that RB966982 amplifies its glycolytic capacity far more dramatically upon infection. This high flux glycolytic state likely fuels the energetic and biosynthetic demands of an effective defense, while CTC9001’s weaker induction may limit its ability to mount robust resistance. In a smaller scale, these same metabolic changes can be observed by DEGs under the 6-phosphofrutokinase GO term (GO:0003872) across all contrasts (sup. Fig 3).

#### 4.1.7. Classical growth x defense tradeoffs hallmark the transcriptional profiles of CTC9001 and RB966982 when challenged by *P. zeae*

To reconcile the patterns from our GO term network analysis, we generated two heatmaps showing the top 30 DEGs for the MCM complex (GO:0042555) and photosynthesis (GO:0015979). In the CTC_inoc vs. RB_inoc comparison, genes linked to cell-division were significantly induced in the CTC variety upon inoculation. While, in the RB_control vs. RB_inoc comparison, photosynthesis pathways were notably repressed in nematode infected roots. The MCM complex heatmap (sup. Fig 4), shows 30 out of 42 DEGs belonging to this GO term. Among these top 30, 28 are highly induced (Z-scores > 1) in the CTC variety upon inoculation, conversely, the same 28 genes are highly repressed in RB variety (Z-scores < 1), suggesting a pause in the host’s cell division, and thus, vegetative growth upon nematode inoculation. Complementary to the data provided by the MCM complex analysis across the treatments, the photosynthesis heatmap shows 30 out of 128 DEG’s under this GO term, in which the RB_control shows high basal photosynthetic activity, but represses this machinery in a significant manner upon nematode infection, this is even more notably given that this transcriptome is based in root, and not foliar tissue. These changes highlight distinct transcriptional changes in each variety, while CTC promotes growth and cell division upon nematode infection, RB halts its photosynthetic pathways in a robust manner (sup. Fig 5).

#### 4.1.8. Complex lipid metabolism underlines the resistant RB9666982 transcriptional profile

The hierarchical clustering of the top 30 (out of 152) differentially expressed genes (DEGs) annotated under the fatty acid biosynthetic process (GO:0006633) reveals a distinct transcriptional signature between varieties (sup. Fig 6). Within this subset, the expression patterns reflect the broader trend observed across 152 DEGs, highlighting a pronounced basal transcriptional profile associated with lipid metabolism in the RB variety compared to the CTC variety. Additionally, several small clusters of genes exhibit specific induction in the RB variety following nematode inoculation. Overall, these data demonstrate that complex lipid metabolism, specifically the coordinated up-regulation of fatty acid biosynthetic enzymes, is a defining feature of the resistant RB966982 transcriptional profile under pathogen attack (Figure 11). This selective induction likely contributes to the production of lipid-derived signaling compounds (e.g., oxylipins, jasmonates) and membrane remodeling events that are a hallmark of effective defense. Additionally, a lipid metabolism–related gene was observed being expressed in the resistant cultivar RB966982 at high levels regardless of pathogen challenge, whereas in the susceptible CTC9001 line its expression remained at or near the detection limit under both control and inoculated conditions. Specifically, arachidonate 12-lipoxygenase (SoffiXsp0nR570.05Cg177900) showed a higher median and mean expression in RB966982 mock samples than after inoculation, yet overall abundance and variability changed little upon infection (Figure 12). These results may indicate that RB966982 relies on a constitutively active oxylipin pathway, driven by basal 12-lipoxygenase activity, to mount its defense, while the absence of this preformed expression in CTC may contribute to its susceptibility, especially in the context of overall lipid metabolism differences between these two varieties.

**Figure 12.**
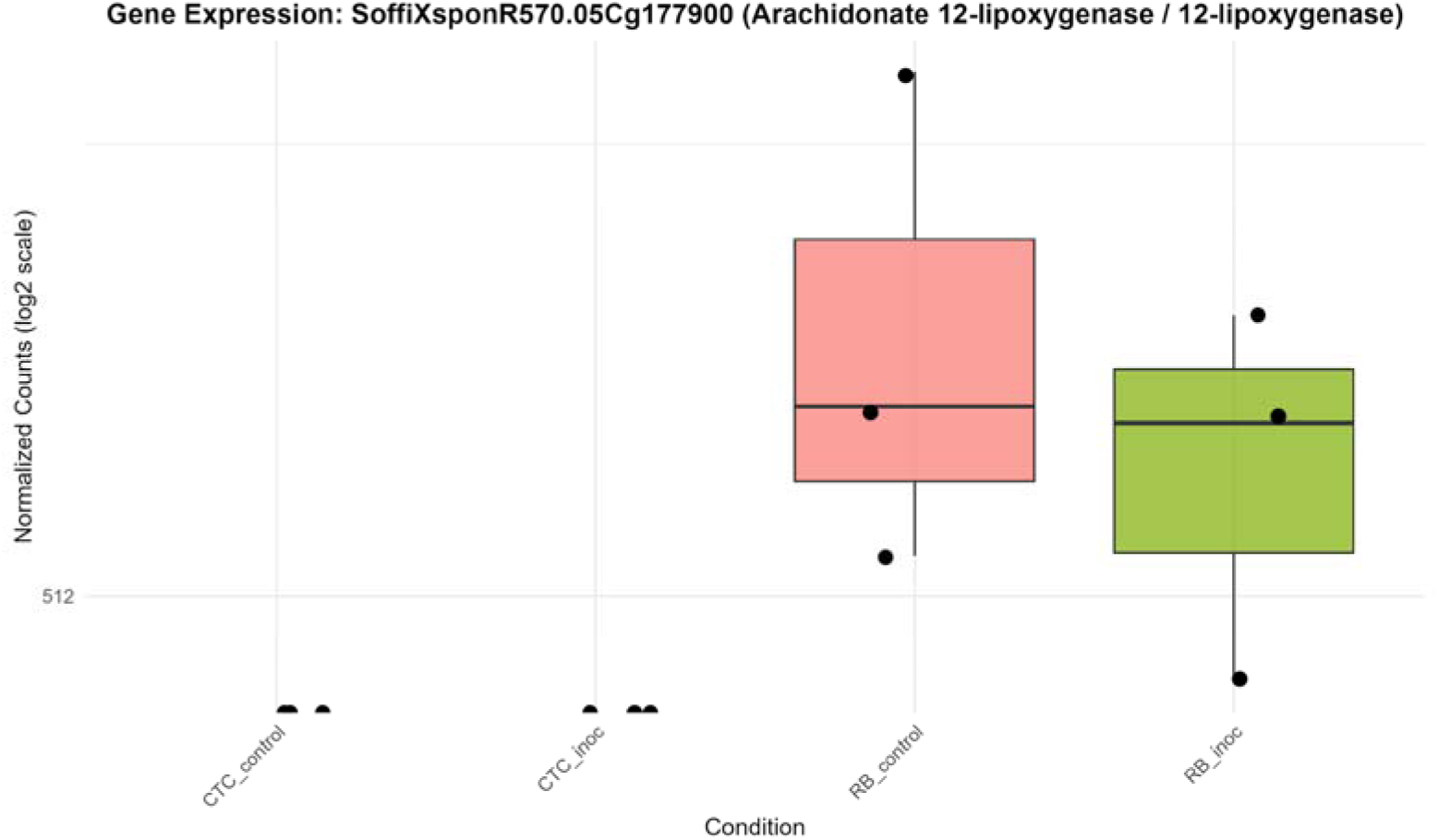
Boxplots illustrating expression patterns (normalized counts) of arachidonate 12-lipoxygenase (SoffiXsp0nR570.05Cg177900) across all contrasts sugarcane varieties under control and inoculated conditions at 15 DAI.

#### 4.1.5. Opposing Transcriptional Regulation of PR1 Homologs in Resistant RB9666982 and Susceptible CTC Cultivars upon Nematode Inoculation

To identify key DEGs, we searched for genes that were differentially expressed in both varieties upon inoculation and then filtered for those whose direction of change was reversed between the inoculated contrasts. In total, 109 genes met these criteria: 39 were significantly up-regulated in CTC-inoculated roots and down-regulated in RB966982-inoculated roots, while the remaining 70 displayed the opposite pattern (Supplementary Material). Notably, within the subset up-regulated in RB966982 upon inoculation, and down-regulated in CTC, we found 12 pathogenesis-related protein 1 (PR1) homologs (Figure 13), suggesting that PR1 induction could be a key component of the resistant cultivar’s defense response.

**Figure 13.**
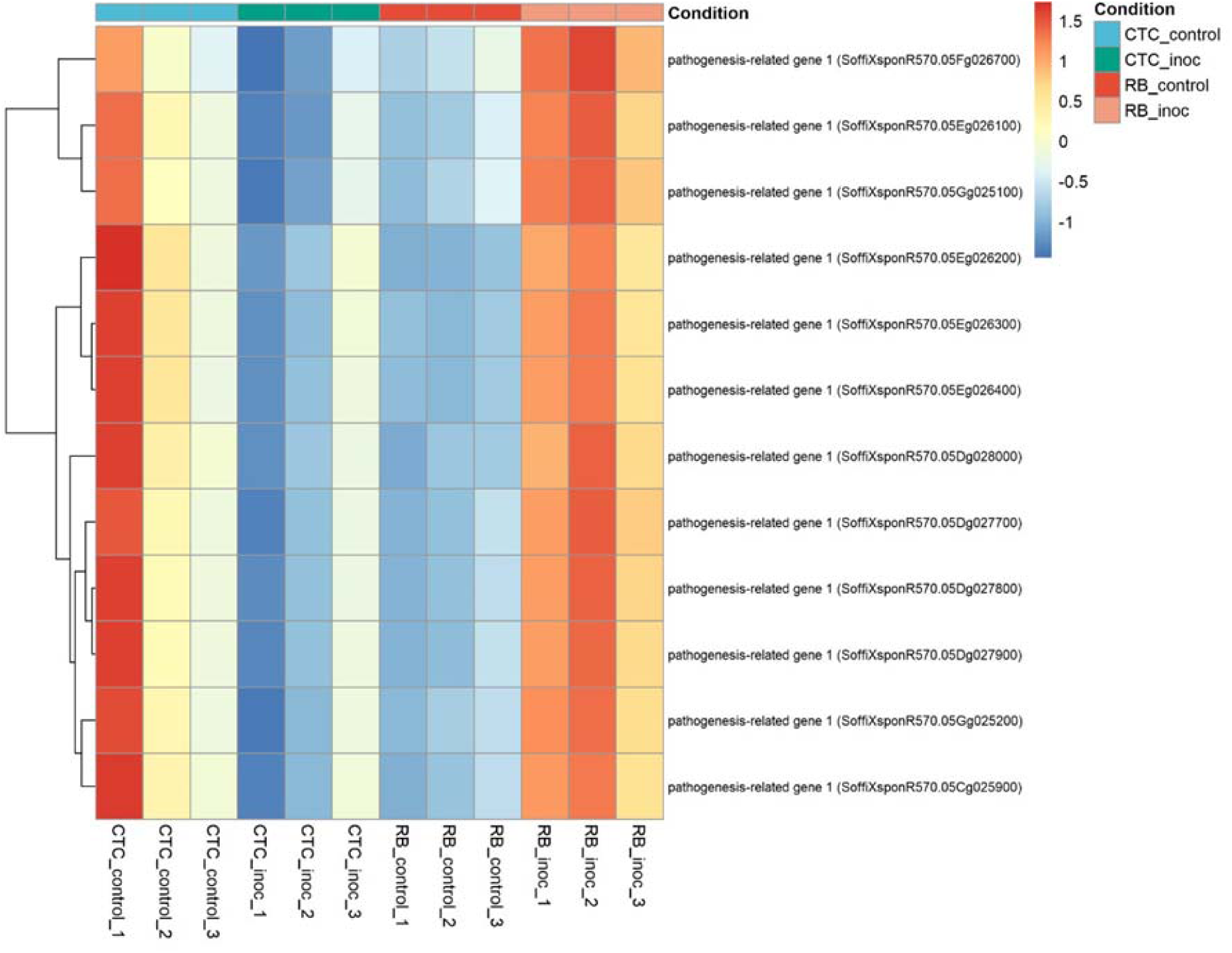
Heatmap representing the normalized expression (Z-score) of 12 pathogenesis-related genes across susceptible (CTC9001) and resistant (RB966982) varieties, comparing control and inoculated conditions at 15 DAI. Color scale indicates relative expression levels ranging from highly induced (red) to strongly repressed (blue). Gene clustering (rows) reflects similarity in expression patterns across samples (columns).

## 5. DISCUSSION

Unveiling the transcriptional profiles underlying resistant and susceptible host responses to root lesion nematodes is a key step in developing strategies to reduce yield losses, particularly those caused by soilborne pathogens. In this sense, the study here presented is the first to conduct a comprehensive investigation into the molecular mechanisms involved in the sugarcane resistance response to *P. zeae*. Differential gene expression analysis and GO term enrichment revealed that both the resistant (RB966928) and susceptible (CTC9001) varieties undergo robust cellular reprogramming 15 days after nematode inoculation, triggering coordinated metabolic changes. Strikingly, the resistant variety exhibited far broader transcriptional adjustments, displaying 2.5 times more differentially expressed genes (DEGs) than its susceptible counterpart, most of which were upregulated. While published transcriptome studies of *Pratylenchus* remain limited, the expression pattern here observed aligns with a similar study in resistant soybean challenged by *P. brachyurus*, albeit at 8 dai (Lopes-Caitar et al., 2022). Additionally, inter-variety contrasts allowed us to observe substantial differences between CTC9001 and RB966928 even in the absence of the pathogen (mock inoculation control), a difference that would grow even larger upon nematode inoculation. This inter-variety inoculated contrast allowed us not only to observe overall expression profiles changes induced by the nematode presence in each genotype but also identify transcripts that were absent in one variety, and thus, undetectable in their respective CTC/RB (control x inoculated) contrasts. These observations were in line with the PCA analysis, which identified host genotype as the primary determinant of transcriptional profiles. Such patterns are consistent with other transcriptome studies comparing distinct genotypes within the same species, in which the variable of interest, in our case the nematode inoculation, comes as a secondary driving factor of transcriptional change (Richards et al., 2012; Sharma et al., 2025).

Through the GO term enrichment analysis, we were able to observe relevant patterns throughout this pathosystem interactions. The first remarkable feature is that, even though nematode inoculation was a secondary driver of transcriptional changes, it was the primary driver of GO term induction/suppression. All contrasts involving nematode inoculation presented more enriched GO terms than the inter-variety control comparison. This is an indication that the nematode presence (regardless of resistance status), induces a more coordinated transcriptional response in the presence of this biotic stressor (Figure 14).

**Figure 14.**
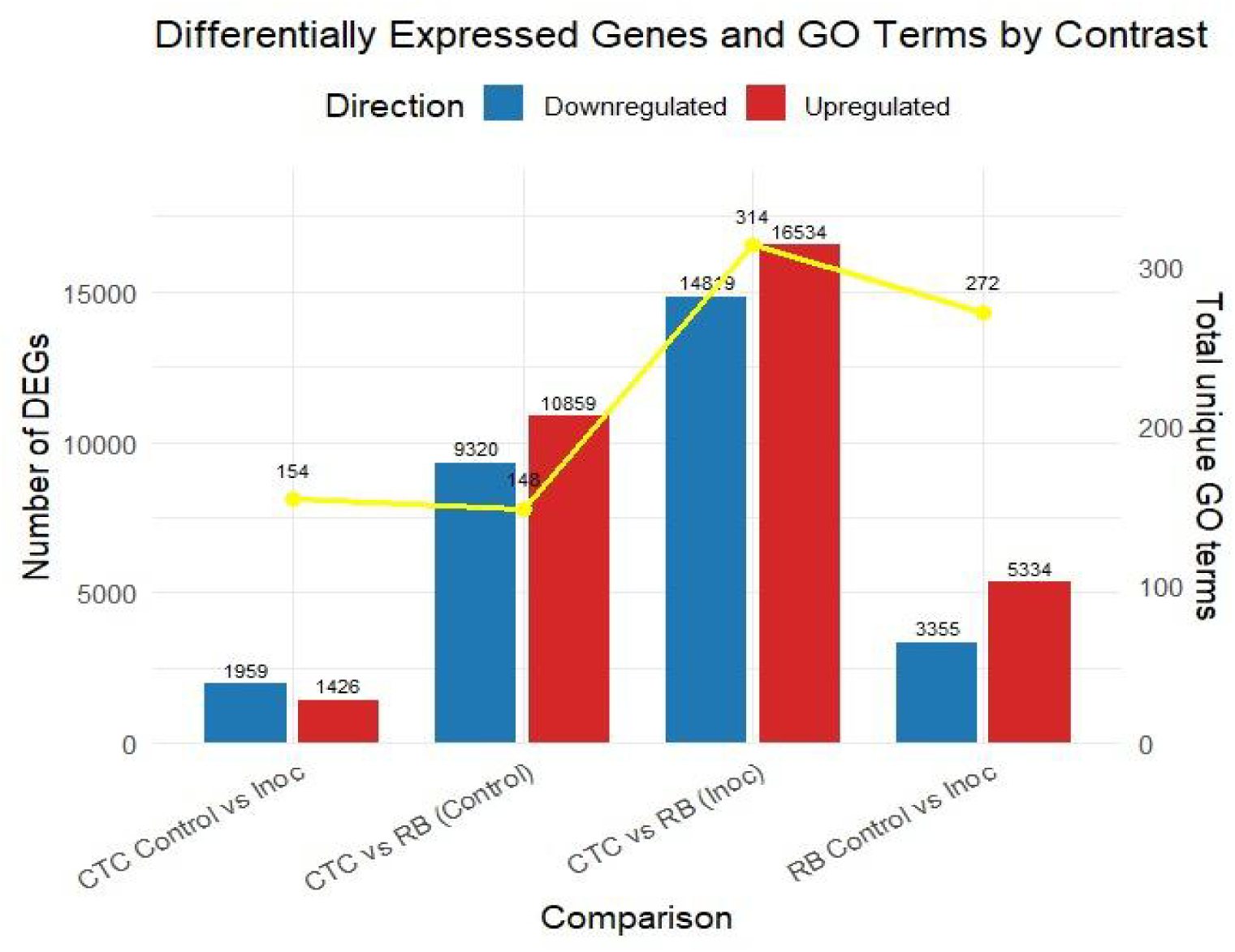
Bar plot displaying the number of significantly upregulated (red) and downregulated (blue) genes across four pairwise comparisons. Differentially expressed genes (DEGs) were identified using a threshold of adjusted *p*-value (padj) ≤ 0.05 and an absolute **log**_₂_ **fold-change of (|log**_₂_**FC|) > 1**. Enriched GO terms (FDR < 0.05) for each contrast are presented as the yellow line in the secondary axis.

The top eight enriched GO terms (FDR < 0.05) in each contrast, grouped by upregulated and downregulated genes revealed consistent biological patterns associated with nematode presence across all inoculated comparisons.

In the CTC_control x CTC_inoc contrast, which marks our susceptible response by the CTC9001 variety (Figure 5), induced dominant terms are marked by a strong network of microtubule motor activity and kinesin complex genes, hallmarking the activity of a synergetic complex responsible for cellular reorganization, and transport of key metabolites, which is in line with an expected defense response (Yun & Kwon, 2017). Notably, cytoskeletal rearrangements terms observed in this contrast have been linked to plant immune response attributed to rapid reorganization of cell structures, transportation of cell compounds, and to reactive oxygen species (ROS) feedback-regulation and salicylic acid (SA) pathway signaling (Matoušková et al., 2014; J. Wang et al., 2022). To further back up this hypothesis of a tentative defense response by the susceptible contrast, we have three enriched GO terms in a hub centered around the 1,3-β-D-glucan metabolism, responsible for the biosynthesis of callose. In a plant-pathogen interaction, the building of callose depositions occurs between the cell membrane and cell wall and consists of repeating glucose units linked by β-1,3 glycosidic bonds, these depositions play a role as a mechanical barrier to delay intracellular access to the pathogen (B. Wang et al., 2022; Y. Wang et al., 2021a). In response to nematode infection, callose deposition is induced as a primary defense mechanism. This response is triggered by pathogen-associated molecular patterns (PAMPs) and pattern-triggered immunity (PTI), which are activated upon recognition of conserved nematode pheromones (e.g., ascarosides) and mechanical disruption of cell walls during *P. zeae* migration and feeding (Chen et al., 2024; Manosalva et al., 2015). An alternative hypothesis to the notion that the susceptible CTC9001 is mounting initial defense responses at 15 DAI can be proposed by observing the prevalent GO terms when the susceptible and resistant inoculate hosts are compared in the inter-variety contrasts CTC_inoc vs. RB_inoc (Figure 7). In this comparison, a complex network of enriched GO terms associated with cellular multiplication can be observed, implying that the CTC9001 variety might be prioritizing cellular multiplication under *P. zeae* attack, and the terms regarding cytoskeletal, microtubule and kinesin activity could in fact be reflecting their role in cell division, rather than defense (Bellelli & Boulton, 2021; Ganguly et al., 2012). In this context even the 1,3-β-D-glucan metabolism could be attributed to cellular division, as callose deposition is a key step in the formation of the cell plate before complete cytokinesis in plant cells (Hong et al., 2001).

The repressed terms of the intra-variety contrast (CTC_control vs CTC_inoc) reveal distinct signatures of nematode parasitism (Figure 5). These downregulated patterns highlight systemic alterations of host processes, including: (1) suppression of nutrient reservoir activity and metal ion homeostasis, (2) dampening of defense responses, and (3) disruption of protein synthesis machinery. First, the halting of nutrient accumulation could be attributed to the fueling of a defense response or the direct build-up of antimicrobial compounds being used from inside the vacuoles (de Souza Cândido et al., 2011). Alternatively, this could indicate that these reserves are being invested towards cellular division, as discussed in the alternative hypothesis to CTC’s susceptibility. Second, the downplay of defense responses are here presented in the repression of the terms defense response to bacterium (GO:0042742) and defense response to fungus (GO:0050832). While not directly related to nematode parasitism, these terms underscore several genes involved in general plant response defense and have been found to be enriched under a successful plant defense against nematodes (Sato et al., 2021), pointing to a possible increased susceptibility in the CTC9001 at 15 DAI. Third, the repression of transcriptional machinery as presented by downregulation of genes involved in the eukaryotic translation elongation factor 1 complex (eEF1) indicates a global reduction in protein synthesis capacity. This downregulation of eEF1 complex genes can play dual role in hampering plant defenses, first by compromising the synthesis of defense related proteins such as pattern recognition receptors (PRR) or pathogenesis related proteins (PRs), and secondly by playing a role in negatively regulating programmed cell death (Son & Park, 2023; S. Wang et al., 2017).

As an overview, CTC9001’s susceptible response profile exhibits two possible characteristic patterns: (1) delayed/attenuated defense activation, or alternatively (2) enhanced cell division during pathogen challenge. In both scenarios, the susceptible host demonstrates simultaneously: a mobilization of nutrient reserves, a suppression of protein synthesis machinery, and the downregulation of defense-related pathways, collectively indicating successful pathogen manipulation of host physiology.

The resistant RB966982 genotype presents a markedly different metabolic profile when challenged by *P. zeae* at 15 DAI (Figure 6). Its defense strategy relies heavily in an integrated biochemical network, including glycolysis intensification to fuel ATP production, phosphofructokinase (PFK) activity surges to drive carbon flux (Yao & Wu, 2016), while malic enzymes and malate dehydrogenase (MDH) maintain a steady pulse of redox power (Gautam et al., 2017). This metabolic interplay seems intended to sustain a carefully balanced influx of ROS signals and oxidoreductase activity, creating physiological conditions for sustained defense activation while maintaining tissue integrity, which is in line with a successful defense response to plant pathogenic nematodes (Meresa et al., 2024; Qiao et al., 2023; Zacheo et al., 1997). The induction of PFKs can be seen in sup. figure 3, where several putative 6-phosphofructokinase (PFK6) genes are up-regulated in the resistant variety after infection. Interestingly, the same heatmap shows a cluster of 5-phosphofructokinase (PFK5) genes with opposite regulation patterns between the susceptible and resistant plants. This is noteworthy because existing studies on PFK5 have only described its function in chloroplasts (Hess et al., 2021; Yoshida & Hisabori, 2021), while our data comes from root tissues. This robust defense response further includes upregulation of thiamine pyrophosphate-binding genes (GO:0031993), consistent with established resistance mechanisms. Thiamine metabolism plays a dual protective role, serving both as an antioxidant system and as a precursor for defense-related metabolite synthesis (Tunc-Ozdemir et al., 2009; Yusof, 2019).

The inter-variety comparison (CTC vs. RB) under both mock-inoculated and nematode-inoculated conditions revealed distinct patterns in basal expression and resistance responses related to lipid metabolism. Notably, RB966982 exhibited induced GO terms associated with *fatty acid biosynthetic process* (GO:0006633, BP) 15 days after inoculation (DAI). Intriguingly, the mock-inoculated RB966982 plants also showed induction of *lipid transport* (GO:0006869, BP) at the same point, suggesting a constitutively higher basal activity in lipid metabolism compared to CTC9001. This constitutive elevation in lipid metabolism observed in RB966982 could play a factor in priming the host for a more robust and rapid defense response when challenged by *P. zeae* (Sarowar et al., 2009). Enhanced fatty acid biosynthesis and transport can lead to increased production of signaling molecules such as jasmonic acid (JA), which is synthesized from linolenic acid via the octadecanoid pathway (Weber, 2002). JA plays a pivotal role in orchestrating defense responses against a broad spectrum of nematodes, by activating the expression of defense-related genes and facilitating cross-talk with other hormonal pathways like salicylic acid (SA) (Meresa et al., 2024; Zacheo et al., 1997). Therefore, the induction of lipid metabolism in RB966982 might not only contribute to the biosynthesis of key signaling molecules but also to the integration of broader network of hormonal interactions and stress signaling pathways that collectively enhance the plant’s resistance to nematode infection. Additionally, lipids play key roles in regulating ROS formation, another defense strategy employed by the resistant variety, as discussed before (Seth et al., 2024).

Supporting the hypothesis of a JA-mediated response, the 12-lipoxygenase (12-LOX) (SoffiXsp0nR570.05Cg177900) showed cultivar-specific expression, being detected only in the resistant line (Figure 12). In plants, the LOX family of enzymes catalyzes the oxygenation of polyunsaturated fatty acids to form 12-hydroperoxy derivatives, in a pathway that ultimately leads to the synthesis of JA (Viswanath et al., 2020). In monocots such as maize, the 9-LOX (ZmLOX3) pathway is required for effective JA-dependent defenses against root-parasitic nematodes, with ZmLOX3 mutants showing increased susceptibility, illustrating the role of LOXs in nematode resistance (Gao et al., 2008). Curiously, besides its role in defense signaling, JA has a direct nematocidal effect on *P. zeae* when exposed in vitro as demonstrated by Gavin et al. (2013). Therefore, data suggest that in the resistant variety, this basal elevation of 12-LOX expression might contribute to the role of metabolic pre-arming and integrating lipid signaling with hormonal crosstalk, especially between JA and salicylic acid (SA), establishing a robust multilayered defense network that the susceptible variety lacks.

Hormonal crosstalk is an essential part of plant defense, with the classical antagonistic model between JA and SA, emerging as a base of discussion. In this interplay, JA coordinates responses to necrotrophs and chewing insects, while SA primarily defends against biotrophs and hemibiotrophs (Caarls et al., 2015; Li et al., 2019). Although this paradigm holds true for many pathosystems, plant-parasitic nematodes appear to challenge this conventional view. As demonstrated by Manosalva et al. (2015) in *Arabidopsis*, the aforementioned nematode MAMP, ascarosides, elicits a PTI response similar to classical MAMPs such as flagellin, however, recognition of the nematode leads to the simultaneous accumulation of SA and JA, leading to defense patterns observed in both pathways. Previously, it was shown that enhanced JA signaling contributes to RB966982’s resistance to *P. zeae*. Intriguingly, in this resistant variety we also observed strong up-regulation of 12 pathogenesis-related (PR1) genes, classic markers of SA signaling (S. Ali et al., 2018), upon inoculation, whereas the susceptible line fails to induce PR1 (Figure 13). This dual JA–SA activation suggests that RB966982 not only mounts an oxylipin-driven defense but also engages SA-mediated responses to reinforce its barrier against nematode feeding. The role of a SA-mediated response is further supported by the robust oxi/redox metabolism observed in the RB966982, one of the hallmarks of this hormonal defense pathway (Herrera-Vásquez et al., 2015).

As another layer of defense, cell wall modifications emerge as a crucial component in the resistance response evidenced by our induced GO terms (RB_control vs. RB_inoc). While the susceptible CTC9001 showed strong correlation between nematode presence and increased callose deposition precursors (1,3-β-D-glucans) as seen in sup. Fig 1, the resistant RB966982 instead upregulated transcription of cell-wall modifying enzymes from the xyloglucan fucosyltransferases family. This distinction is reflected in the contrasting GO term profiles: 1,3-β-D-glucan-related activities (GO:0003843, GO:0051274, GO:0017011) dominated the susceptible response, while galactoside 2-alpha-L-fucosyltransferase activity (GO:0015031) characterized the resistant genotype. These patterns are clearly visible in the DEG heatmaps for each contrast (sup. Fig 2). To distinguish these cell-wall modifying strategies, it is necessary to highlight their differences. As mentioned before, callose deposition is a fast defense response after pattern recognition receptor (PRR) activation, forming a physical line of defense. However, callose deposits act as a temporary barrier that gets broken down once the stress is over (Li et al., 2023; Y. Wang et al., 2021b). Conversely, fucosyltransferases (FUT) provide a more basal and durable form of resistance by catalyzing α-1,2-fucosylation of cell-wall polysaccharides, most notably the addition of L-fucose to xyloglucan side chains and pectic components, which reinforces wall architecture, impedes pathogen-secreted hydrolases, and optimizes glycosylation of immune receptors, thereby establishing a long-lasting structural fortification throughout the cell wall. A vast body of literature exists on molecular and physiological defense responses to sedentary plant-parasitic nematodes (e.g., *Meloidogyne*, *Heterodera*). In these interactions, callose appears as a crucial component of the host’s defense arsenal. Notably, callose deposits are observed surrounding feeding sites (giant cells or syncytia) reprogrammed by these nematodes, likely in an attempt to block nutrient flow to the parasitized tissue (M. A. Ali et al., 2013; Holbein et al., 2016). This defense strategy was detailed in a study by Hofmann et al., (2010), in which higher callose content in the plasmodesmata surrounding feeding sites was linked to smaller syncytia and giant cells, impacting *Meloidogyne incognita* and *Heterodera schachtii* reproduction on *Arabidopsis thaliana* and *Nicotiana tabacum*, respectively. This might suggest that the susceptible CTC9001 is indeed manifesting a broad general response to nematode recognition. However, this strategy seems unfruitful, likely due to the mode of parasitism explored by *P. zeae*, which is migratory for nematodes of the genus *Pratylenchus*. This feeding strategy involves continuous migration through the root cortex, with brief feeding interactions lasting anywhere from 5 minutes (short feeding) to 2 hours (long feeding), and unlike sedentary nematodes, this relationship is non-permanent (Zunke, 1990). This divergence in cell-wall defense strategy between the susceptible CTC9001 and the resistant RB966982 seems at least partially responsible for the observed phenotypes, in which a callose focused defense appears unable to hinder nematode development, while the permanent fucosylation of cell wall components at a tissue level poses a challenge to *P. zeae* parasitism.

Corroborating the fact that CTC9001 mounts a broad defense response rather than a focused one, Figure 9 shows that the vast majority of upregulated RGAs were observed in the susceptible interaction. While this pattern might appear unexpected, such an “overshoot” of defense genes in susceptible interactions is not uncommon and may reflect a generalist, ineffective response (Schenk et al., 2000), in this case, as a contrast to the more coordinated defense seen in RB966982. This pattern of defense-related genes being broadly expressed in a susceptible response is observed in a study by Vieira et al. (2019), in which the resistant alfalfa cultivar MNGRN-16 expressed far fewer defense-related transcripts than the susceptible cultivar Baker upon *Pratylenchus penetrans* infection.

Focusing on the up-regulated RGAs in the resistant response, two interesting patterns emerged, especially represented by the NB-ARC domain genes SoffiXsponR560.05Dg100600, SoffiXsponR560.09Eg112400 and SoffiXsponR560.09oe1a063500, all up-regulated in the inter-variety inoculated contrasts (Figure 10). The first pattern, characterized by SoffiXsponR560.05Dg100600, consists of distinct basal expression levels that respond positively to nematode inoculation. The second pattern, represented by the other two genes, shows three distinct expression profiles: (1) near absence in CTC9001, (2) high variability in mock-treated plants, and (3) strong induction in RB966982 following infection. This dual strategy, appears to be combining a pre-existing reservoir of RGA transcripts with an infection-triggered amplification, resembling the phenomenon of defense priming, in which elevated basal levels of key immune components poise the plant for faster and stronger activation upon challenge (Chen et al., 2024; Conrath et al., 2015). In the same line, a review by Tsuda & Katagiri, (2010) highlights that robust effector-triggered immunity depends on early orchestration of both pattern-triggered and RGA–mediated networks, with pre-existing transcript pools enabling a more rapid hypersensitive response. By contrast, CTC9001 must mobilize the same RGAs that from near-zero expression upon nematode attack, leading to a delayed, broad-spectrum “overshoot” that often arrives too late to stop infection. Together, these patterns suggest that RB966982’s superior resistance stems from layering basal readiness with inducible reinforcement, ensuring both immediate effector recognition and sustained signaling throughout the immune response.

## 6. CONCLUSION

This study provides the first comprehensive transcriptomic analysis of sugarcane’s response to *Pratylenchus zeae* infection, comparing resistant (RB966928) and susceptible (CTC9001) varieties. Key findings reveal that the resistant genotype employs a multifaceted defense strategy, characterized by robust metabolic reprogramming, enhanced lipid signaling (particularly JA-mediated pathways), and coordinated hormonal crosstalk between JA and SA. Additionally, cell wall reinforcement via fucosyltransferases activity and pre-armed defense priming through elevated basal expression of resistance gene analogs (RGAs) appears to play an essential role to RB966982’s resistance. In contrast, the susceptible CTC9001 variety exhibits either a delayed/attenuated defense response or a shift toward cellular proliferation, accompanied by broad but ineffective activation of defense-related genes, such as those involved in callose deposition. In synthesis, this work not only advances our understanding of sugarcane-*P. zeae* interactions but also opens the door to innovative strategies for nematode resistance, including marker-assisted breeding, targeted genetic modifications, and the development of stable genetic resistance approaches.

## Declarations

### Ethics approval and consent to participate

Not applicable.

### Consent for publications

Not applicable.

### Availability of data and materials

The datasets generated during the current study are available in the NCBI BioProject database (http://www.ncbi.nlm.nih.gov/bioproject/) under accession number PRJNA1309697, accession IDs for the samples are provided in the supplementary file 2. Code for the reproduction of the analyses within this paper is available on GitHub at https://github.com/pedroconfort/pzeaesugarcaneinteraction.

### Competing interests

The authors declare that they have no competing interests.

### Funding

This study was supported by the Fundação de Amparo à Pesquisa do Estado de São Paulo (FAPESP–2022/03962). Aperfeiçoamento de Pessoal de Nível Superior - Brasil (CAPES) is reponsible for the support in the form of salary for author PMSC through the PIPD post-doctoral scholarship.

## Supporting information

Accessions table

## Acknowledgments

We thank Elaine Vidotto Batista (Genomics Group Lab, ESALQ/USP). **Authors’ contributions:** CBM-V conceived the study and edited the manuscript. PMSC analyzed the data and wrote the manuscript. TGS and JF conducted the experiments. SC contributed with plant materials.

## Supplementary Information

**Additional file 1 (.docx):** Document containing the supplementary figures 1 to 7.

**Additional file 2 (.docx):** Table of accessions by sample.

## Notes

### Competing Interest Statement

The authors have declared no competing interest.

