## Supplementary Figures for "Multilayered Defense Responses in Sugarcane Against *Pratylenchus zeae* Revealed by Comparative Transcriptomics"

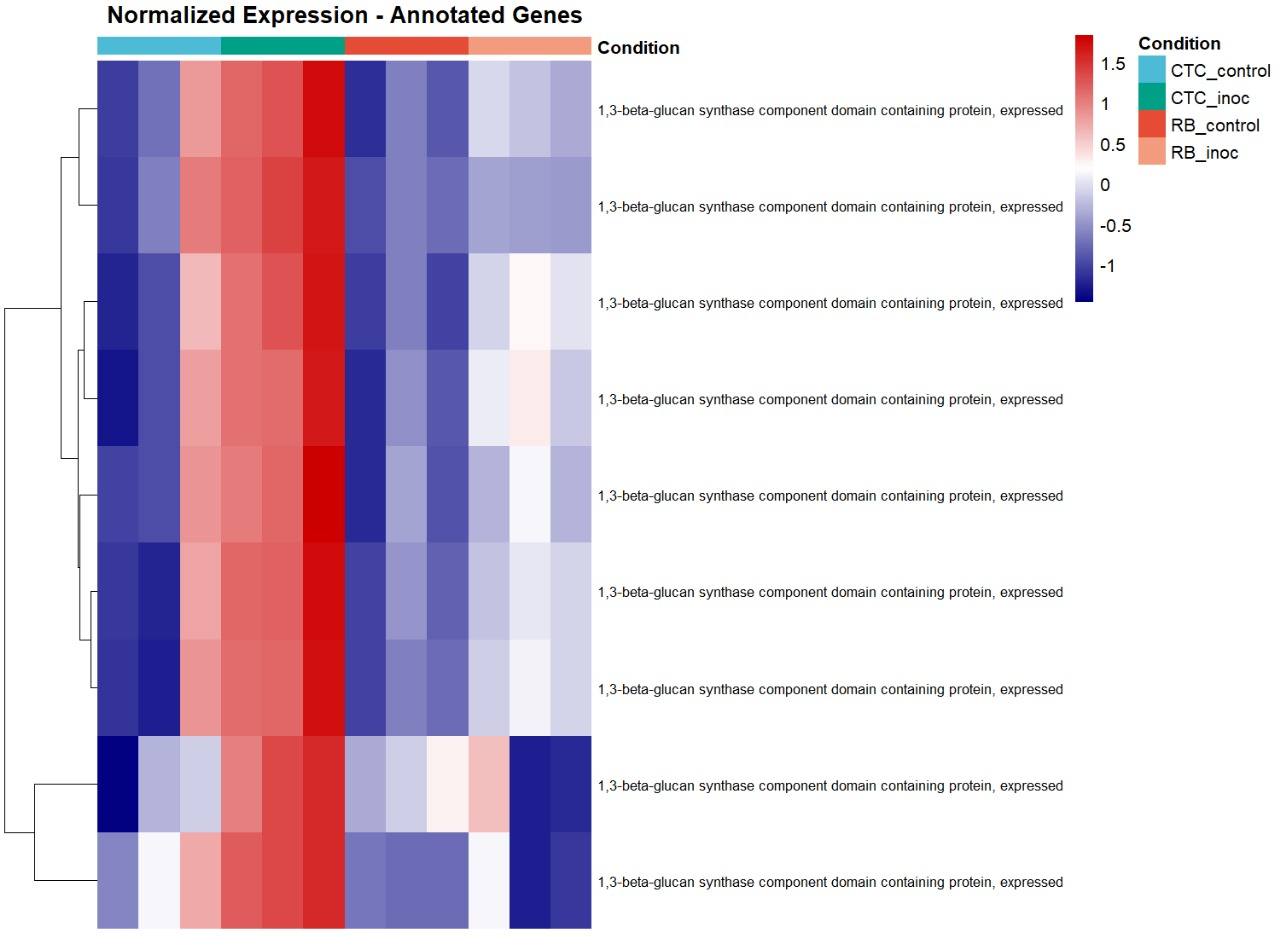
Supplementary figure 1: Normalized expression (Z-score) of annotated 1,3-beta-glucan synthase genes across susceptible (CTC9001) and resistant (RB966982) varieties, comparing control and inoculated conditions at 15 DAI.

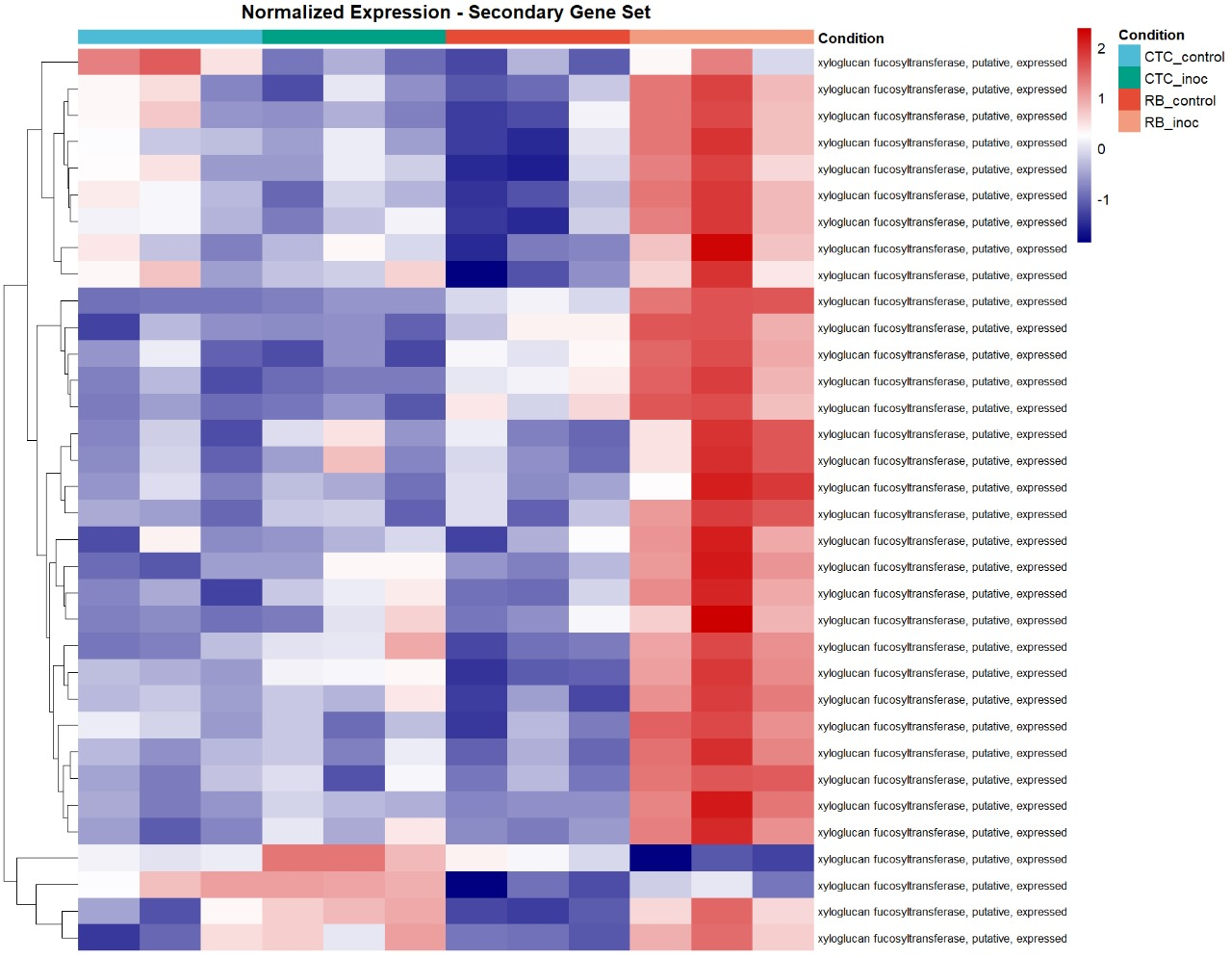

Suplementary figure 2: Heatmap representing normalized expression (Z-score) of annotated xyloglucan fucosyltransferases- genes across susceptible (CTC9001) and resistant (RB966982) varieties, comparing control and inoculated conditions at 15 DAI. Color scale indicates relative expression levels ranging from highly induced (red) to strongly repressed (blue). Gene clustering (rows) reflects similarity in expression patterns across samples (columns).

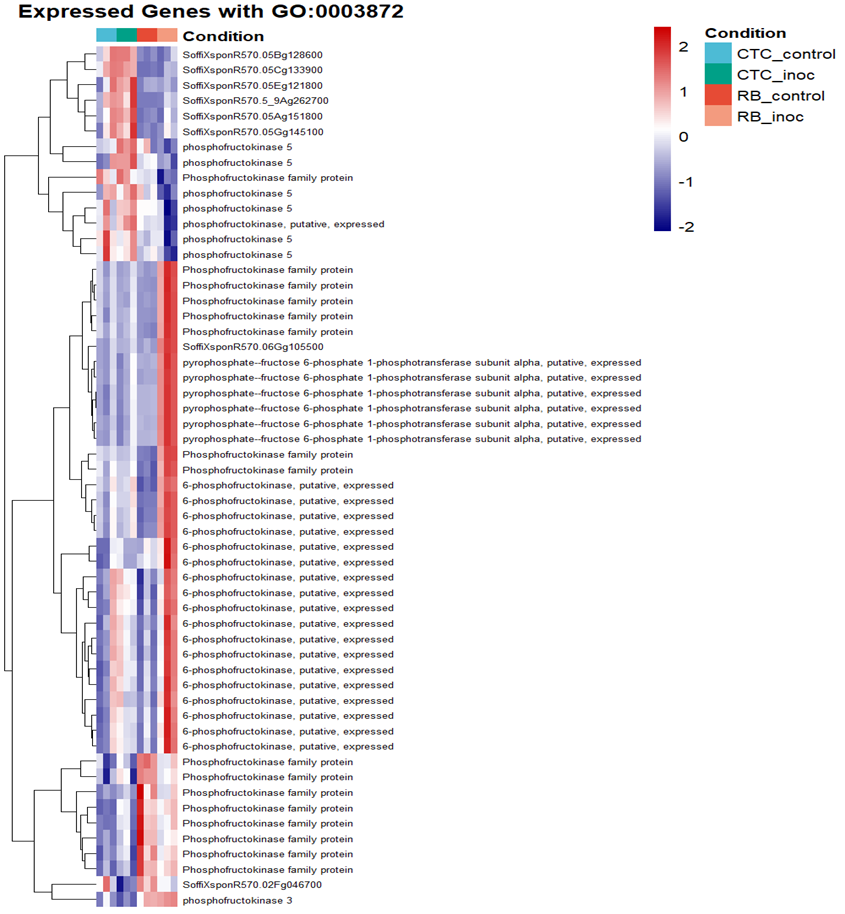

Supplementary figure 3: Heatmap illustrating Z-score–normalized expression of all genes annotated 6-phosphofructokinase activity (GO:0003872) across susceptible (CTC9001) and resistant (RB966982) varieties under both control and inoculation conditions at 15 DAI. Color scale indicates relative expression levels ranging from highly induced (red) to strongly repressed (blue). Gene clustering (rows) reflects similarity in expression patterns across samples (columns).

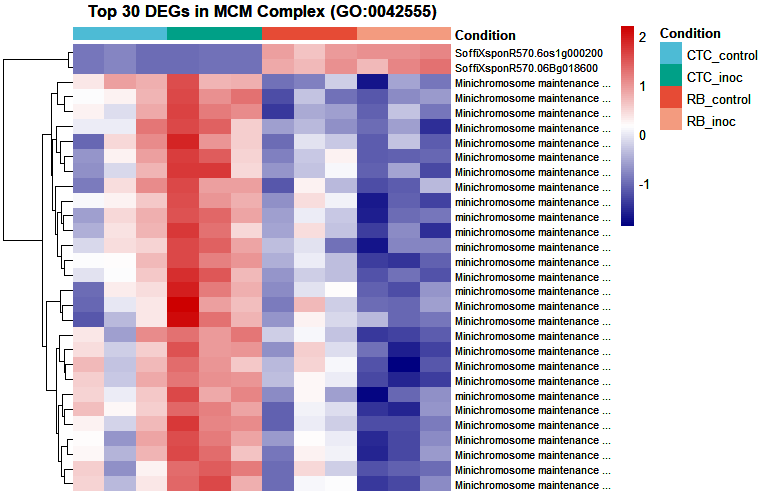

Supplementary figure 4: Heatmap illustrating Z-score–normalized expression of the top 30 DEGs annotated to the MCM complex (GO:0042555) across susceptible (CTC9001) and resistant (RB966982) varieties under both control and inoculation conditions at 15 DAI. Color scale indicates relative expression levels ranging from highly induced (red) to strongly repressed (blue). Gene clustering (rows) reflects similarity in expression patterns across samples (columns).

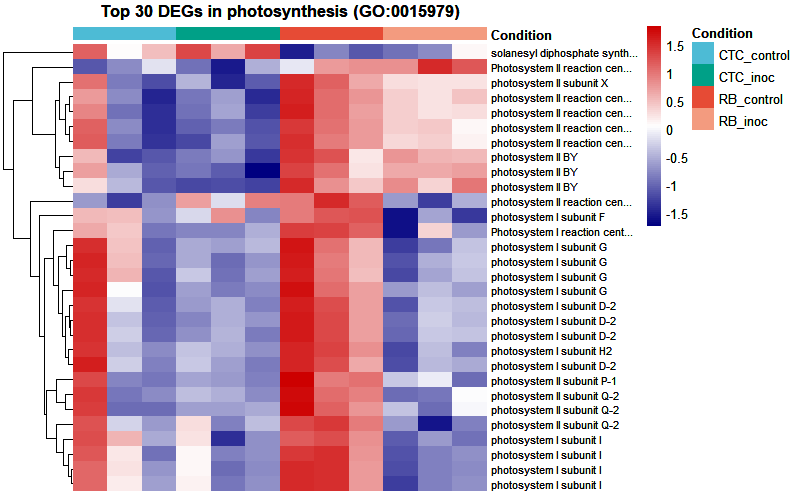

Supplementary figure 5: Heatmap illustrating Z-score–normalized expression of the top 30 DEGs annotated to Photosynthesis (GO:0015979) across susceptible (CTC9001) and resistant (RB966982) varieties under both control and inoculation conditions at 15 DAI. Color scale indicates relative expression levels ranging from highly induced (red) to strongly repressed (blue). Gene clustering (rows) reflects similarity in expression patterns across samples (columns).

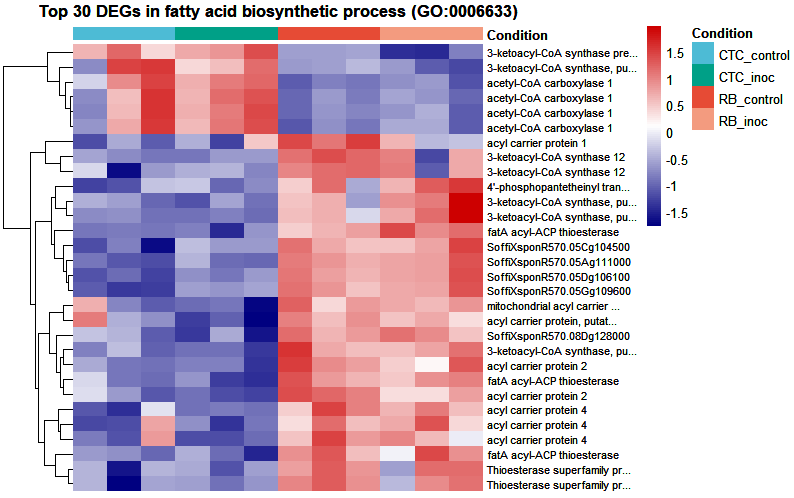

Supplementary figure 6: Heatmap illustrating Z-score–normalized expression of the top 30 DEGs annotated to fatty acid biosynthetic process (GO:0006633) across susceptible (CTC9001) and resistant (RB966982) varieties under both control and inoculation conditions at 15 DAI. Color scale indicates relative expression levels ranging from highly induced (red) to strongly repressed (blue). Gene clustering (rows) reflects similarity in expression patterns across samples (columns).

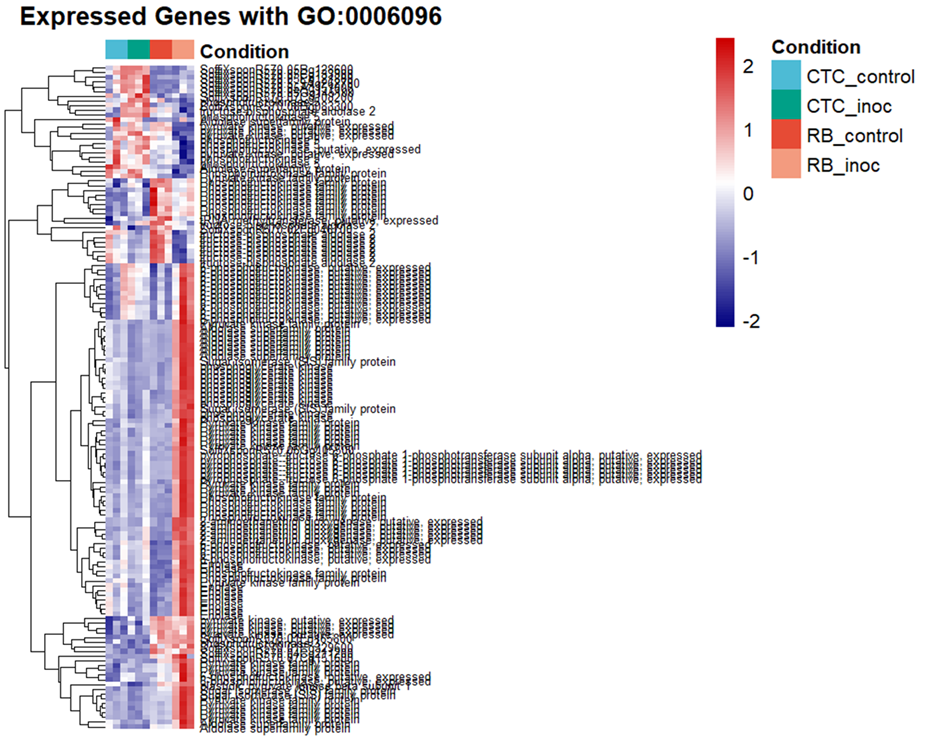

Supplementary figure 7: Heatmap illustrating Z-score–normalized expression of the top 30 DEGs annotated to glycolytic-process (GO:0006696) across susceptible (CTC9001) and resistant (RB966982) varieties under both control and inoculation conditions at 15 DAI. Color scale indicates relative expression levels ranging from highly induced (red) to strongly repressed (blue). Gene clustering (rows) reflects similarity in expression patterns across samples (columns).
