## Supplementary material for "Multilayered Defense Responses in Sugarcane Against *Pratylenchus zeae* Revealed by Comparative Transcriptomics": Accessions table

| Accession | Sample | SPUID | Organism | ID | Cultivar | BioProject |
| --- | --- | --- | --- | --- | --- | --- |
| SAMN50751030 | CTCCR1_S124_L001_R1_001 | CTCCR1_S124_L001_R1_001 | Saccharum hybrid cultivar | 128810 | CTC9001 | PRJNA1309697 |
| SAMN50751031 | CTCCR2_S125_L001_R1_001 | CTCCR2_S125_L001_R1_001 | Saccharum hybrid cultivar | 128810 | CTC9001 | PRJNA1309697 |
| SAMN50751032 | CTCCR3_S126_L001_R1_001 | CTCCR3_S126_L001_R1_001 | Saccharum hybrid cultivar | 128810 | CTC9001 | PRJNA1309697 |
| SAMN50751033 | CTCInoR1_S121_L001_R1_001 | CTCInoR1_S121_L001_R1_001 | Saccharum hybrid cultivar | 128810 | CTC9001 | PRJNA1309697 |
| SAMN50751034 | CTCInoR2_S122_L001_R1_001 | CTCInoR2_S122_L001_R1_001 | Saccharum hybrid cultivar | 128810 | CTC9001 | PRJNA1309697 |
| SAMN50751035 | CTCInoR3_S123_L001_R1_001 | CTCInoR3_S123_L001_R1_001 | Saccharum hybrid cultivar | 128810 | CTC9001 | PRJNA1309697 |
| SAMN50751036 | RBCR1_S118_L001_R1_001 | RBCR1_S118_L001_R1_001 | Saccharum hybrid cultivar | 128810 | RB966982 | PRJNA1309697 |
| SAMN50751037 | RBCR2_S119_L001_R1_001 | RBCR2_S119_L001_R1_001 | Saccharum hybrid cultivar | 128810 | RB966982 | PRJNA1309697 |
| SAMN50751038 | RBCR3_S120_L001_R1_001 | RBCR3_S120_L001_R1_001 | Saccharum hybrid cultivar | 128810 | RB966982 | PRJNA1309697 |
| SAMN50751039 | RBInoR1_S115_L001_R1_001 | RBInoR1_S115_L001_R1_001 | Saccharum hybrid cultivar | 128810 | RB966982 | PRJNA1309697 |
| SAMN50751040 | RBInoR2_S116_L001_R1_001 | RBInoR2_S116_L001_R1_001 | Saccharum hybrid cultivar | 128810 | RB966982 | PRJNA1309697 |
| SAMN50751041 | RBInoR3_S117_L001_R1_001 | RBInoR3_S117_L001_R1_001 | Saccharum hybrid cultivar | 128810 | RB966982 | PRJNA1309697 |

<https://www.ncbi.nlm.nih.gov/biosample/50751030>
<https://www.ncbi.nlm.nih.gov/biosample/50751031>
<https://www.ncbi.nlm.nih.gov/biosample/50751032>
<https://www.ncbi.nlm.nih.gov/biosample/50751033>
<https://www.ncbi.nlm.nih.gov/biosample/50751034>
<https://www.ncbi.nlm.nih.gov/biosample/50751035>
<https://www.ncbi.nlm.nih.gov/biosample/50751036>
<https://www.ncbi.nlm.nih.gov/biosample/50751037>
<https://www.ncbi.nlm.nih.gov/biosample/50751038>
<https://www.ncbi.nlm.nih.gov/biosample/50751039>
<https://www.ncbi.nlm.nih.gov/biosample/50751040>
<https://www.ncbi.nlm.nih.gov/biosample/50751041>
